# Pac1/LIS1 promotes an uninhibited conformation of dynein that coordinates its localization and activity

**DOI:** 10.1101/684290

**Authors:** Matthew G. Marzo, Jacqueline M. Griswold, Steven M. Markus

## Abstract

Cytoplasmic dynein is a minus end-directed microtubule motor that transports myriad cargos in various cell types and contexts. How dynein is regulated to perform all these activities with a high degree of spatial and temporal precision is unclear. Recent studies have revealed that human dynein-1 and dynein-2 can be regulated by a mechanism of autoinhibition, whereby intermolecular contacts limit motor activity. Whether this autoinhibitory mechanism is conserved throughout evolution, whether it can be affected by extrinsic factors, and its precise role in regulating cellular dynein activity remain unknown. Here, we use a combination of negative stain EM, single molecule motility assays, genetic, and cell biological techniques to show that the autoinhibitory conformation is conserved in budding yeast, and it plays an important role in coordinating dynein localization and function in cells. Moreover, we find that the Lissencephaly-related protein, LIS1 (Pac1 in yeast) plays an important role in regulating this autoinhibitory conformation of dynein. Specifically, our studies demonstrate that rather than inhibiting dynein motility, Pac1/LIS1 promotes dynein activity by stabilizing the uninhibited conformation, which ensures appropriate localization and activity of dynein in cells.

## INTRODUCTION

Cytoplasmic dynein is an enormous minus end-directed microtubule motor complex that transports numerous cargoes. At first glance, this motor seems exceedingly complex in terms of its architecture, size, and reliance on accessories and regulators for proper activity. For instance, processive single molecule motility of human dynein – itself comprised of 4 to 6 subunits – requires the 11 subunit dynactin complex in addition to an adaptor that links them together^1, 2^. Although yeast dynein does not require dynactin for *in vitro* single molecule motility^3^, it does require this complex for *in vivo* activity^4, 5^. Recent studies have yielded invaluable insight into the underlying reasons for the complexity of the dynein motor. For instance, the reliance on adaptors (*e.g.*, BicD2, Spindly, Hook3^1, 2^) to link dynein to dynactin ensures that cytoplasmic dynein-1 – which effects motility of numerous and varied cargoes throughout the cell cycle^6^ – and dynactin are linked together at the right place (and presumably time) for appropriate motility. Additionally, recent studies have revealed that dynactin helps to orient the motor domains in a parallel manner that is conducive for motility^7^, thus revealing the mechanistic basis for dynein’s reliance on this large complex. Thus, the complexity of this molecular motor ensures that cargoes are transported to their target destinations in accordance with the needs of the cell.

In addition to its regulation by extrinsic factors, several studies have demonstrated that human dynein-1 and dynein-2 can also be auto-regulated by intra-complex interactions. Specifically, intermolecular interactions between the motor domains have been shown to stabilize an autoinhibited conformation of human dynein called the phi particle (named for its resemblance to the Greek letter)^7–10^. In the case of dynein-2 (responsible for intraflagellar transport), the phi particle conformation – which has been observed in its native context^10^ – reduces its velocity, ATPase activity and microtubule landing rate^9^. Similarly, the autoinhibited dynein-1 conformation has been shown to reduce its microtubule landing rate and motility properties^7, 11^. Moreover, unlike dynein-2 which is not regulated by dynactin^12^, uninhibited dynein-1 mutants interact more readily with dynactin and the adaptor BicD2^7^.

Although it is well established that human dynein adopts the autoinhibited phi particle conformation (both dynein-1 and dynein-2), it is unclear if this conformational state is evolutionarily conserved. Yeast dynein is of particular interest due to two notable *in vitro* discrepancies with human dynein. In particular, unlike human dynein, yeast dynein is processive in single molecule assays without the need for other factors, such as dynactin^3^. The second notable feature that distinguishes yeast dynein is its apparent ability to interact with dynactin in the absence of any additional factors (*i.e.*, adaptors)^13^. The reasons for these differences are unclear, but one possibility is that yeast dynein does not adopt the autoinhibited phi particle conformation, which could potentially account for its ability to walk in the absence of dynactin. This is supported by studies showing that artificially separating the motor domains of human dynein-1 with a rigid linker (thus preventing intermolecular contacts) is sufficient to convert it to a processive motor^11^.

In addition to dynactin, the Lissencephaly protein LIS1 is another important effector of dynein activity that is required for numerous dynein functions in cells^14, 15^. These include promoting dynein recruitment to various cellular sites^16, 17^, and assisting in dynein transport functions, including nuclear migration in neurons^18, 19^, and high-load vesicular transport^20–23^. However, the mechanism by which LIS1 affects dynein, or dynein-dynactin activity remains controversial. For instance, *in vitro* studies have shown that LIS1 can either reduce^23–25^ (for dynein alone) or increase dynein velocity (in the context of intact dynein-dynactin-BicD2 complexes)^25, 26^. In addition to promoting *in vitro* force production by dynein^23^, studies have also shown that LIS1 can help in the initiation of dynein-dynactin-BicD2 motility from the plus ends of dynamic microtubules^25, 27^.

Studies with the budding yeast homolog of LIS1 – Pac1 – have shown that it reduces the velocity of dynein motility^28–31^, presumably by uncoupling the ATPase cycle from the conformational changes in the motor and microtubule-binding domains that elicit microtubule release^29, 30^. Thus, the precise role for Pac1/LIS1 in dynein function remains confounded by these contrasting results. Although a role for Pac1/LIS1 in regulating the autoinhibited conformation has not yet been reported, two studies found that LIS1 can indeed promote dynein-dynactin interaction^32, 33^, which is an expected consequence of relieving dynein autoinhibition^7^.

Here, we set out to address the question of whether yeast dynein adopts an autoinhibited phi particle conformation, and if so, what role it plays in regulating *in vitro* and *in vivo* dynein activity. Our recent findings suggested the potential for yeast dynein to adopt such a conformation^34^. Specifically, we found that engineering a neurological disease-correlated mutation into yeast dynein leads to increased run lengths of single molecules of dynein, and a localization pattern in cells that is suggestive of an enhanced dynein-dynactin interaction. This mutation was within the linker domain – the mechanical element responsible for the powerstroke^35^ – at a residue that was recently shown to be important for maintenance of the phi particle conformation of human dynein^7^. Here, we use a combination of single particle analysis (by negative stain EM), single molecule motility assays, genetic, and cell biological approaches to show that yeast dynein indeed adopts a phi particle conformation that restricts its *in vitro* processivity, and coordinates it localization and activity in cells. Moreover, we find that Pac1 is an important regulator of the phi particle conformation. In particular, we found that rather than inhibiting dynein motility, Pac1 promotes its activity by stabilizing the ‘open’, uninhibited conformational state. Our findings help explain recent observations with LIS1^25, 26^, and support a model whereby Pac1/LIS1 is a key effector of dynein autoinhibition.

## RESULTS

### Yeast dynein adopts an autoinhibited ‘phi’ particle conformation

We sought to determine whether yeast dynein adopts an autoinhibited conformation (the ‘phi’ particle^8^). To this end, we developed a strategy to isolate biochemical quantities of the intact yeast dynein complex that are of sufficient purity for single particle analysis by negative stain electron microscopy. The yeast dynein complex is comprised of light (Dyn2), light-intermediate (Dyn3), intermediate (Pac11), and heavy chains (Dyn1)^36^. We generated a polycistronic plasmid containing all four dynein complex subunits each flanked by a strong, galactose-inducible promoter (*GAL1p*) on the 5’ end, and a synthetic transcriptional terminator (T_synth3_^37^) on the 3’ end. We included a tandem affinity tag (8His-ZZ, or “HZZ”) followed by either a SNAP or HALO tag on the N-terminus of Dyn1 for purification and fluorescent labeling of the complex, respectively. In addition to these four gene cassettes, the plasmid also contains a *URA3* cassette that provides a sequence for recombination-based genomic integration (into the native *ura3-1* allele) and a selectable marker (see Fig. 1A). Cells with the plasmid integrated into their genome were grown in galactose-containing media, and the dynein complex was subsequently isolated from cell lysates using either tandem nickel/IgG affinity, or IgG affinity alone. We estimate the yield of the overexpressed complex to be at least 50-fold greater than the non-overexpressed complex. Single molecule motility assays confirmed the activity of the overexpressed complex to be nearly identical to the non-overexpressed complex (Fig. 1B; also see Fig. 4B and 5). Importantly, the increased yield permitted us to isolate the complex to a high degree of purity using size exclusion chromatography, which revealed an elution profile nearly identical to the human dynein complex isolated from insect cells (Fig. 1C).

**Figure 1.**
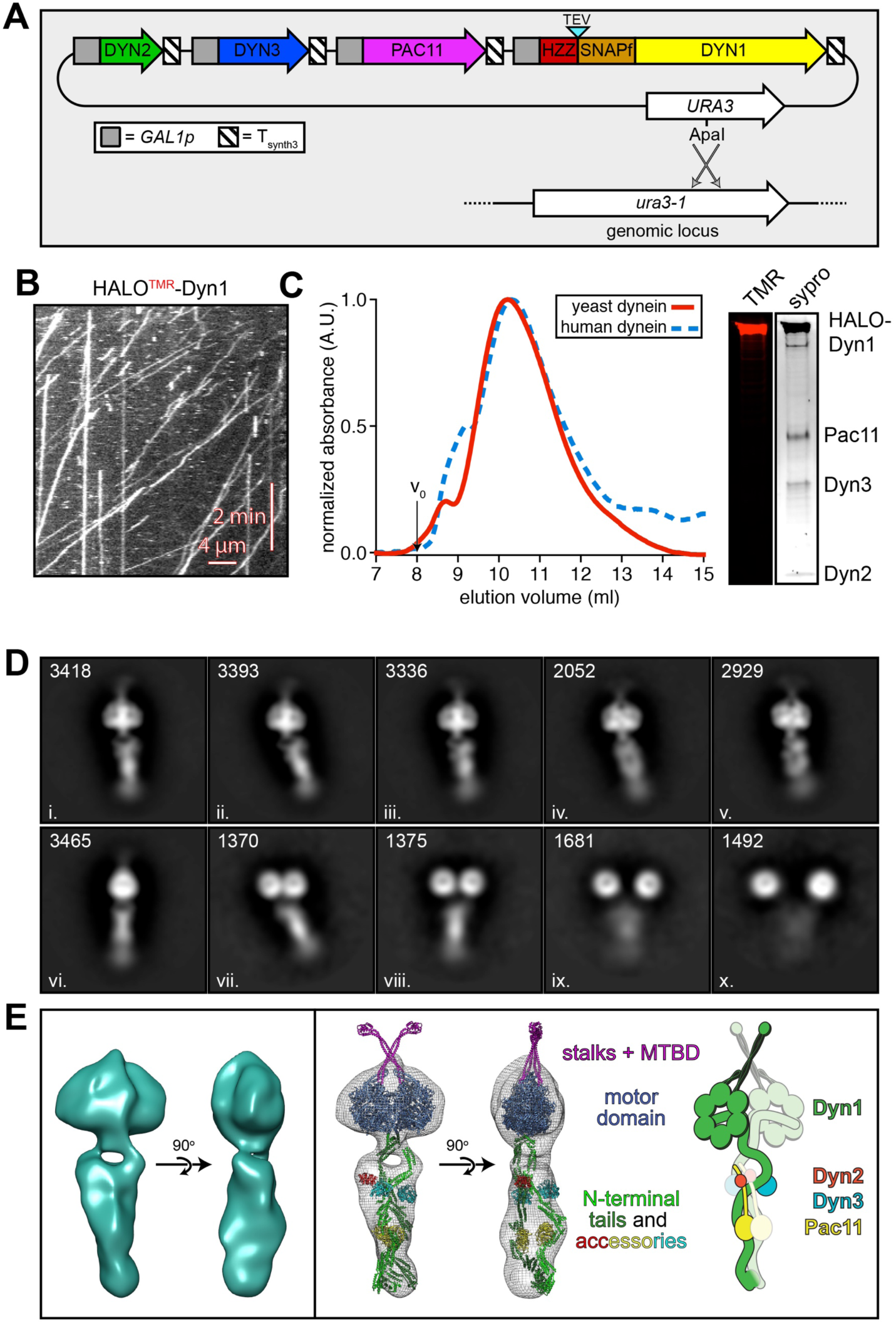
The yeast dynein complex adopts an autoinhibited phi particle conformation. (A) Schematic of the polycistronic plasmid used to produce the intact yeast dynein complex. Restriction digest with ApaI (cuts within *URA3* gene) targets the plasmid for homologous recombination into the *ura3-1* locus as depicted. (B) Representative kymograph depicting single molecule motility of the purified overexpressed yeast dynein complex. (C) Elution profiles of yeast and human dynein complexes from Superose 6 resin (left), and scans of the same polyacrylamide gel depicting fluorescently labeled Dyn1 (via HaloTag-TMR) and the entire complex (via Sypro Ruby staining; right). (D) Representative negative stain EM class averages of the intact yeast dynein complex. Number of particles used to generate each class indicated in each panel. Classes i – vi depict dynein in the autoinhibited, phi particle conformation, whereas vii – x depict dynein in various open, uninhibited states. (E) 3D models of dynein in the autoinhibited state generated from 2D class averages with (right) and without (left) a high resolution 3D structure of human dynein-1 in the phi particle conformation (pdb 5NVU) manually docked into it. Note that the structures of the two tail domains have been slightly rotated with respect to the motor domains to better fit the 3D model, and that the structures of both TcTEX and Robl have been eliminated due to their absence from the yeast dynein complex. Also see Video S1.

With the high purity dynein complex in hand, we used negative stain electron microscopy to obtain the first ever high magnification view of the intact yeast dynein complex. This revealed the presence of several conformational states, including those in an open, uninhibited state (Fig. S1A, green arrow), and those in an apparent phi particle conformation (Fig. S1A, red arrows), with the large majority being in the latter state (Fig. S1B). Reference-free 2D class averages provided images that appear strikingly similar to the intact human dynein-1 complex^1, 7^, and to an artificially dimerized motor domain truncation of dynein-2 in a phi particle conformation^9^ (Fig. 1D). Specifically, the N-terminal tail domains – which exhibit flexibility with respect to the motor domains – appear to be twisted around one another, which we confirmed by generating three-dimensional reconstructions (Fig. 1E, left, and Video S1). Intermolecular contacts appear to extend to the motor domains and the coiled-coil stalks, which connect the AAA ring to the microtubule-binding domains (MTBDs). Much like the human dynein-1 and dynein-2 structures, the stalks cross each other in an “X”-like configuration in a manner that seems contrary to motility. We confirmed the high degree of similarity between the human and yeast dynein-1 phi particle conformations by manually docking a high resolution cryo-EM structure of human dynein (pdb 5NVU^7^) into our 3D model (Fig. 1E, right, and Video S1).

Of note, previous observations of an artificially-dimerized (via glutathione S-transferase, GST), truncated yeast dynein motor domain fragment (lacking the N-terminal tail domain) revealed a lack of phi particle conformations. This includes observations by negative stain EM^28^, and within the crystal lattice of the crystalized motor domain fragment^38, 39^. Thus, in contrast to the dynein-2 isoform of dynein, for which the motor domain is sufficient to adopt the phi particle conformation (as apparent in the crystal lattice and by negative stain EM^9, 40^), yeast dynein requires the tail domain for assembly of this autoinhibited conformation. It is interesting to note that several of our class averages appear to depict a conformation in which the motor domains are closely apposed, but unbound, and the tails appear to be wrapped around one another (Fig. 1D; classes vii and viii). Taken together, this suggests that contact points within the tail domain provide important contacts that likely stabilize the autoinhibited conformation.

### Disruption of the autoinhibited conformation leads to increased single molecule processivity

We previously found that a disease-correlated amino acid substitution within the linker domain – the mechanical element responsible for the powerstroke – leads to an increase in single molecule run length^34^. This residue (K1475) is equivalent to one known to stabilize the autoinhibited conformation of human dynein (R1567)^7^. If the increased processivity is a consequence of disrupted phi particle formation, then we reasoned that mutations at other potential phi particle interfaces would also lead to increased run lengths. A high resolution cryo-EM structure of the human dynein phi particle identified four distinct surfaces that comprise the inter-molecular interface (linker-linker, linker-AAA4, AAA5-AAA5, and stalk-stalk; Fig. 2A)^7^. Given the apparent similarities between the yeast and human dynein phi particles (see Fig. 1E and Video S1), we wondered whether the intermolecular contact points were conserved, and if so, what effect disrupting the phi particle has on dynein motility.

**Figure 2.**
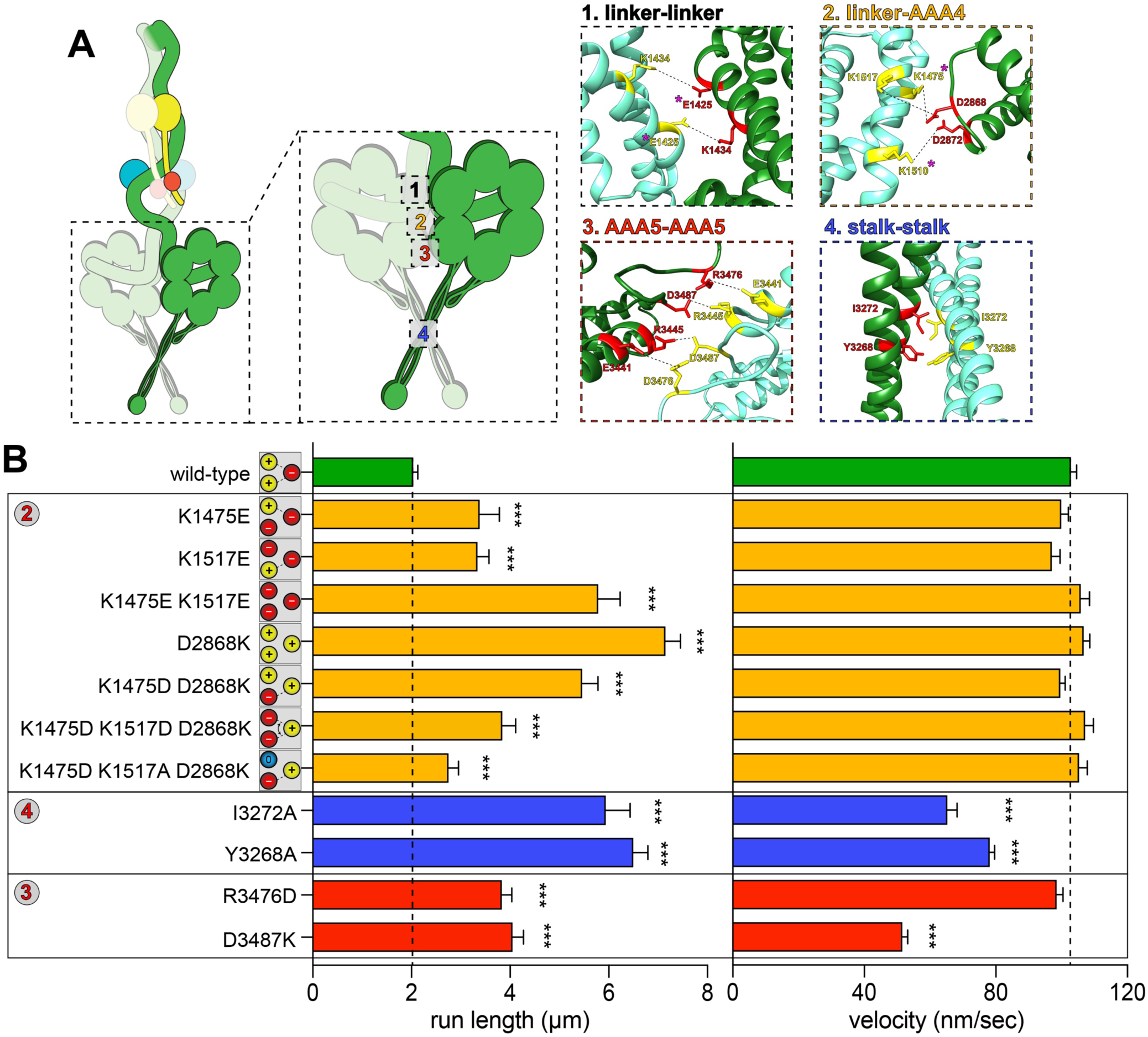
Disrupting phi particle contact points extends single molecule run lengths. (A) Cartoon depicting four predicted intermolecular contact surfaces within the motor domains that stabilize the phi particle conformation. Four insets show respective regions of yeast dynein modeled into human dynein phi particle structure. Structural models were generated using one-to-one threading of the yeast *DYN1* sequence into 5NVU^7^ on the Phyre2 server^72^. Residues with magenta asterisks are mutated in patients suffering from neurological disease^62–64^ (see Discussion). (B) Single molecule run length (from fitting of raw data to one-phase decay) and velocity values for wild-type and mutant dyneins with phi particle disrupting mutations (at surfaces 2, 3 and 4, as indicated). Cartoons along vertical axis depict electrostatic interactions (or lack thereof) among residues 1517, 1475 (left circles) and 2868 (right circle) at linker-AAA4 surface. Note that the degree of processivity enhancement is inversely proportional to the number of charge interactions at this surface. Error bars indicate standard error (between 150 - 528 motors from at least two independent experiments were analyzed for each). Statistical significance was determined using a Mann-Whitney test (for run length) or with an unpaired Welch’s t test (two-tailed; for velocity; ***, p ≤ 0.0001). Also note that we generated and tested the motility of two other point mutants at interface 3, E3441K and R3445D, both of which were inactive in single molecule assays (not shown).

Sequence alignment and homology modeling of pertinent regions of Dyn1 (those at the intermolecular interface) into the high resolution cryo-EM structure of the human dynein phi particle (pdb 5NVU^7^) revealed a high degree of conservation at the four intermolecular surfaces (Fig. 2A). We focused initially on the linker-AAA4 interface (Fig. 2A, surface 2), which is presumably stabilized in part by electrostatic interactions between negatively charged D2868, and positively charged K1475 and K1517. We found that substituting either positively charged residue (K1475 or K1517) with a negatively charged residue (glutamate) resulted in similar increases in single molecule processivity (from 2.0 µm to 3.4 µm and 3.3 µm, respectively; p < 0.0001), while eliminating both (K1475E K1517E) led to an even greater increase in run length (to 5.8 µm; Fig. 2B). Consistent with these residues’ role in stabilizing the linker-AAA4 interface, substituting the negatively charged D2868 for a positively charged one (D2868K) led to an increase in run length that was statistically indistinguishable from the K1475E K1517E double mutant (7.2 µm; p = 0.246). Interestingly, we were able to reduce these run length values by substituting back residues that would be predicted to replace the broken electrostatic pairing (K1475D D2868K; K1475D K1517A D2868K; or, K1475D K1517D D2868K). These results indicate that the linker-AAA4 interface is indeed important for assembly of the autoinhibited conformation of yeast dynein, which attenuates single molecule processivity of the intact complex.

We next wondered if other predicted interfaces affect formation of the autoinhibited conformation, and what role they play in affecting dynein motility. Consistent with the apparent interaction surface observed in our 2D averages within the stalk (Fig. 2A, surface 4; see Fig. 1D, classes i - v), substitution of either Y3268 or I3272 with an alanine led to an increase in run length comparable to those mutants lacking all electrostatic contacts at the linker-AAA4 interface (Fig. 2B). Unlike the linker-AAA4 interface mutants, both Y3268A and I3272A exhibited reductions in velocity values (from 102.8 nm/sec to 73.2 nm/sec for Y3268A, or 78.4 nm/sec for I3272A; p < 0.0001). This could be due to disrupted kinetics of helix sliding in the coiled coil stalk (*i.e.*, changes in the heptad registry), which is responsible for communicating nucleotide-dependent conformational changes within the motor domain to the microtubule-binding domain^40–42^. Finally, charge reversal substitutions at either R3476 (to an aspartate) or D3487 (to a lysine) at the AAA5-AAA5 interface (Fig. 2A, surface 3) also led to increases in run length (from 2.0 µm to 3.8 and 4.0 µm, respectively; p < 0.0001) comparable to the single charge substitution mutants at the linker-AAA4 interface.

Although the R3476D mutant exhibited normal velocity, the D3487K substitution reduced dynein velocity to approximately half that of wild-type. Taken together, these findings confirm the conserved nature of the intermolecular contact points that stabilize the autoinhibited state of yeast dynein. They also indicate that the ability to adopt the phi particle conformation is sufficient to severely limit the processivity of the yeast dynein complex, which, in the absence of phi particle formation, can achieve run lengths that match that of the human dynein-dynactin-BicD2 complex (7.2 µm for dynein^D2868K^ versus 5 - 9 µm for human dynein-dynactin-BicD2)^1, 2, 43^. It is interesting to note that the minimal, GST-dimerized dynein fragment exhibits run lengths (1.6 µm; see Fig S5C) much lower than that achievable by the phi particle disrupting mutants (as high as 7.2 µm, or 4.5-fold higher), in spite of this fragment not adopting the phi particle conformation. This indicates that the native tail domain permits an arrangement of the motor domains that is much more conducive to processive motility than the GST moiety.

### Dynein autoinhibition restricts cortical localization

Although preventing human dynein from adopting the autoinhibited conformation by mutagenesis had no apparent effect on processivity, it did in fact lead to a significant enhancement in the ability of dynein to interact with dynactin and the adaptor BicD2^7^. Previous studies have shown that dynactin is required for localization of dynein to cortical Num1 receptor sites, but not to microtubule plus ends^4, 31, 44^ (Fig. 3A). Moreover, in instances when dynactin interaction with dynein is enhanced^45, 46^, the number of dynein molecules found at cortical sites increases^46^, and dynein offloading to the cell cortex becomes apparent from live cell imaging^31^. In light of the limiting nature of dynactin at microtubule plus ends (1 dynactin complex for every 3 dynein complexes at a plus end)^46^, these observations suggest that interaction with dynactin is a limiting step in the delivery of dynein to cortical Num1 sites. Thus, if disruption of the autoinhibited conformation of yeast dynein leads to enhanced dynactin interaction, then we expect to see an increased frequency of dynein cortical localization.

**Figure 3.**
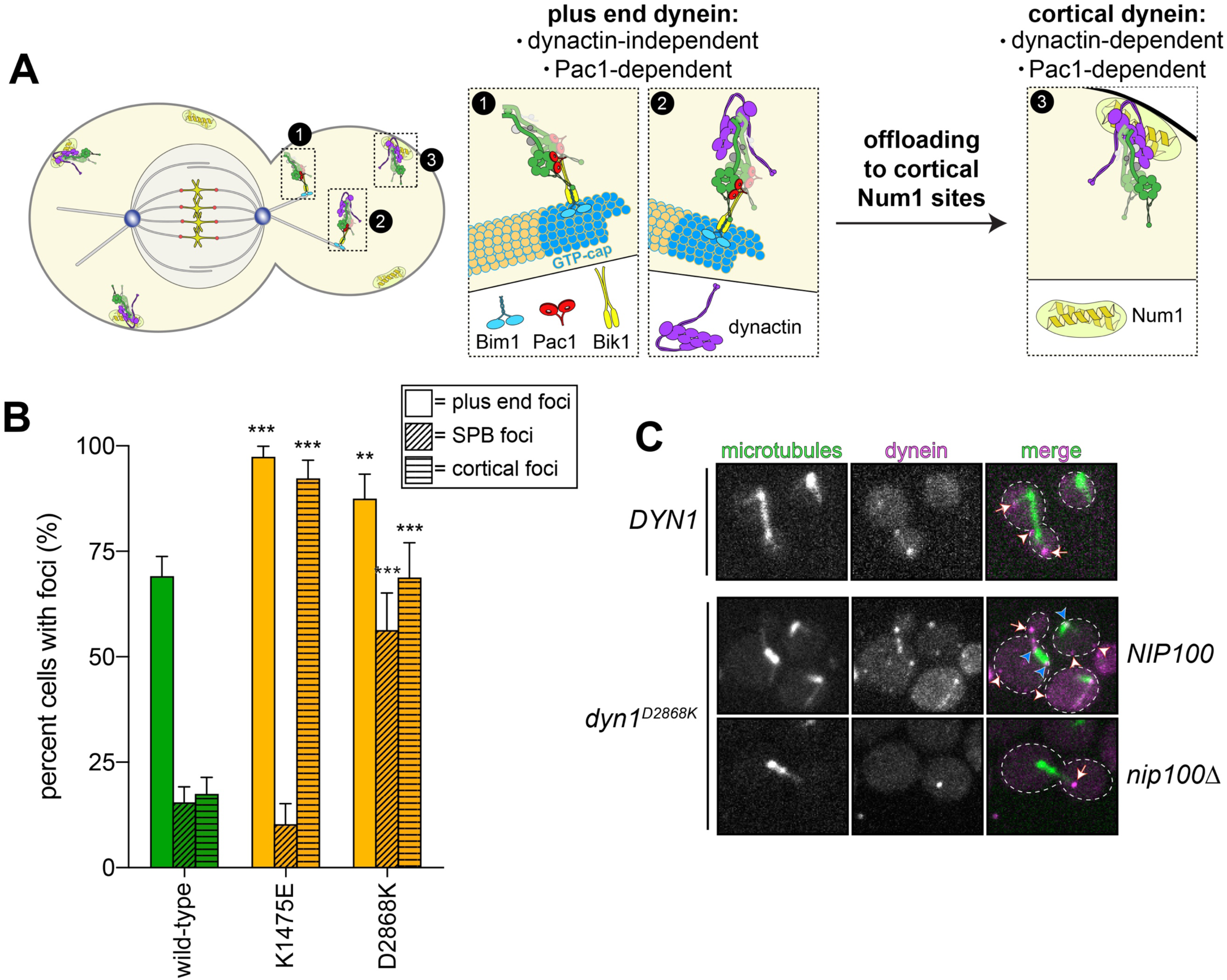
The autoinhibited conformation restricts plus end and cortical localization of dynein. (A) Cartoon depicting the two main sites of dynein localization (microtubule plus ends, and cell cortex), and the molecular requirements for each. Dynein plus end localization (1) requires Bik1^57^ and Pac1^44^, with Bim1 potentially playing some role in this process, but does not require dynactin^4^. Rather, dynactin plus end localization (2) relies on dynein^4^. Subsequent to plus end targeting, dynein-dynactin complexes are offloaded to cortical Num1 sites^31, 73^ (3). (B) Plot depicting the frequency of plus end, SPB (spindle pole body) and cortical targeting for wild-type and mutant Dyn1 (n ≥ 32 mitotic cells for each). Error bars indicate standard error of proportion. Statistical significance was determined by calculating Z scores (see Methods). (C) Representative images of wild-type or mutant dynein (D2868K) localizing in otherwise wild-type or *nip100*Δ (dynactin component) cells. Note the lack of cortical localization of Dyn1^D2868K^ in *nip100*Δ cells (white arrowheads, cortical foci; white arrows, plus end foci; blue arrowheads, SPB foci).

To determine whether this was the case, we compared the localization pattern of wild-type Dyn1-3GFP to that of the K1475E and D2868K mutants, which exhibited modest and strong *in vitro* processivity phenotypes, respectively (see Fig. 2B). Consistent with the notion that disrupting the phi particle promotes interaction with dynactin, we observed a large increase in the number of cells exhibiting cortical dynein foci (Fig. 3B and C). Although D2868K had a stronger processivity phenotype in the single molecule assay, it did not exhibit a stronger cortical localization phenotype than the K1475E mutant. However, we did note that the D2868K cells possessed a higher frequency of dynein foci associated with the spindle pole bodies (SPBs), where the minus ends of microtubules are anchored. Although the relevance of SPB foci is not entirely clear, we previously noted that dynein complexes accumulated at this site when they were activated by overexpression of a Num1 coiled-coil-containing fragment^47^. Thus, the SPB pool of dynein molecules might represent “activated” dynein motors. As the D2868K mutant exhibited a more robust *in vitro* processivity phenotype than K1475E, the former mutation likely results in fewer inhibited dynein molecules in cells. Thus, the increased SPB localization of D2868K is likely a consequence of its increased activity. Finally, we confirmed that dynactin was indeed required for the cortical localization of Dyn1^D2868K^ by imaging its localization in cells lacking the dynactin component, Nip100 (Fig. 3C, *nip100*Δ). These data are consistent with the notion that the phi particle restricts dynein-dynactin interaction, which limits association with cortical Num1 receptor sites.

### Peptide insertion between motor and tail domains ablates phi particle conformation

In addition to an increase in cortical localization, we also noted that the K1475E and D2868K mutants localized to microtubule plus ends to a greater extent than wild-type dynein (see Fig. 3B; p ≤ 0.040). We previously noted that the frequency of plus end localization – which is Pac1-dependent^44^ (see Fig. 3A) – is increased for a truncated dynein motor domain fragment (Dyn1_MOTOR_)^48^, and also for a dynein mutant in which a helical linker peptide (helical linker 3, HL3) was inserted between the tail and motor domains (Dyn1_HL3_; see Fig. 4A)^31^. Of note, Dyn1_HL3_ also localizes to the cell cortex to a greater extent than wild-type Dyn1^31^, much like an isolated tail domain fragment (Dyn1_TAIL_)^48^. We originally generated the Dyn1_HL3_ mutant to test the hypothesis that the motor domain plays a direct role in restricting the cortical localization capacity of Dyn1_TAIL_^31^. We predicted that insertion of the HL3 peptide would sufficiently separate the tail and motor domains such that the motor domain would no longer be able to block the tail domain’s interaction with cortical Num1 receptors (*i.e.*, by “unmasking” the tail domain; see Fig. 4A, “original model”). Although our hypothesis was indeed supported by localization data^31^, there has been no structural or biochemical evidence to support our proposed mechanism of motor-based inhibition of the tail domain. Thus, we wondered whether the HL3 insertion simply disrupts the phi particle conformation (Fig. 4A, “new model”), which would result in an enhanced interaction between dynein, dynactin and Num1.

**Figure 4.**
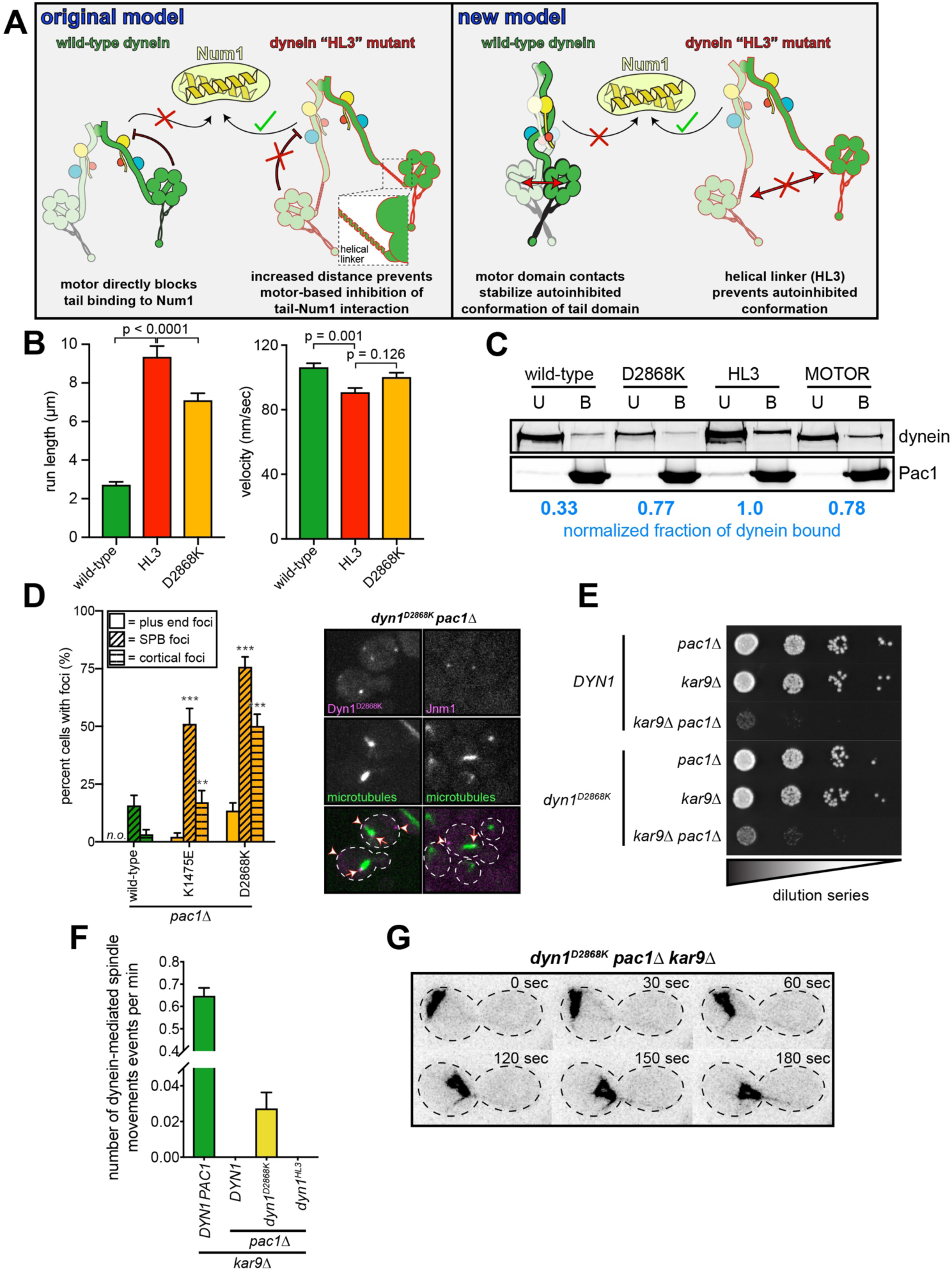
Release of dynein autoinhibition permits Pac1/LIS1-independent localization and function. (A) Cartoons depicting original, and new models accounting for “unmasking” phenotype observed with Dyn1^HL3^ mutant^31^. Wild-type dynein tail domain is unable to associate with Num1 in the absence of plus end-targeting; however, addition of helical linker 3 (HL3) between tail and motor domains permits dynein to associate with Num1 independent of plus end-targeting. Our original model posited that this was a consequence of the motor domain directly precluding the tail domain from contact Num1; however, our new model is that contacts within the motor domain stabilize the phi particle conformation, in which the tail domains are in a twisted state that is unable to interact with Num1. In this latter model, insertion of HL3 prevents the adoption of the phi particle conformation. (B) Single molecule run length (from fitting of raw data to one-phase decay) and velocity values for wild-type and indicated mutant dyneins, as indicated, purified using plasmid-integration strategy described in Figure 1A (n ≥ 224 motors for each, from at least two independent experiments; error bars indicate standard error). Statistical significance was determined using a Mann-Whitney test (for run length) or with an unpaired Welch’s t test (two-tailed; for velocity). (C) Bead binding assay illustrating increased affinity of Pac1 for “open” dyneins (dynein^D2868K^ and GST-dynein_MOTOR_). Purified dyneins were incubated with Pac1-FLAG-SNAP-decorated beads, and the bound (“B”) and unbound (“U”) fractions were resolved by SDS-PAGE. The normalized, relative bound and unbound fractions were determined by measuring band intensities. (D) Plot depicting the fraction of cells with indicated mutant or wild-type Dyn1 foci in *pac1*Δ cells (n ≥ 34 mitotic cells for each; “*n.o.*”, none observed; error bars indicate standard error of proportion). Representative fluorescence images depicting the presence of cortical dynein (Dyn1) and dynactin (Jnm1) in *dyn1^D2868K^* cells (arrowheads, cortical foci; arrows, SPB foci). Statistical significance was determined by calculating Z scores (**, p = 0.011; ***, p ≤ 0.0001). (E) Serial dilutions of cells with indicated genotype were plated on rich media (YPA supplemented with 2% glucose) and grown at 30°C for 2 days (as shown) or 4 days (see Fig. S3A). Note the partial rescue of cell viability in *kar9Δ pac1Δ dyn1^D2868K^* cells as compared to *DYN1 kar9Δ pac1*Δ cells. (F) Plot depicting number of dynein-mediated spindle movements per cell per minute in hydroxyurea (HU)-arrested cells (all of which are *kar9*Δ; see Methods; n ≥ 32 HU arrested cells for each). (G) Representative time-lapse fluorescence images of a hydroxyurea (HU)-arrested *dyn1^D2868K^ pac1Δ kar9*Δ cell exhibiting a dynein-mediated spindle movement.

Single molecule analysis of Dyn1_HL3_ revealed that this mutant exhibits run lengths that exceed all other phi particle disrupting mutants (9.3 µm; p ≤ 0.0001), suggesting that HL3 peptide insertion indeed disrupts phi particle formation more than any of the single point mutants (Fig. 4B). Since we expressed the HL3 mutant complex using the strategy described in Figure 1, we compared its motility to similarly isolated wild-type and D2868K complexes. We noted that although dynein^HL3^ and dynein^D2868K^ exhibited significantly longer run lengths than wild-type dynein (p < 0.0001), the overexpressed wild-type dynein complex exhibited slightly longer runs than the non-overexpressed complex (2.0 µm versus 2.7 µm, p = 0.016; see Fig. S2A). The same was not true for dynein^D2868K^ (7.2 µm for native, versus 7.3 µm for overexpressed; p = 0.9488). This suggests that the phi particle conformation of the overexpressed dynein complex is more labile than the non-overexpressed complex. Consistent with this notion, the run length of the overexpressed wild-type complex – but not the one expressed from native promoters – increased over time to similar levels as the D2868K mutant, even when stored at −80°C (Fig. S2; also see below). By comparison, we only observed a minor increase over time for the non-overexpressed wild-type complex, and for the overexpressed dynein^D2868K^. This is similar to observations with human dynein, which needs to be prepared fresh in order to obtain a sufficient proportion of phi particles^7^.

We previously noted that, like the truncated motor domain fragment^48^ (both the monomeric^48^ and artificially dimerized^3^ variants) – which does not adopt the phi particle conformation (see above) – the Dyn1_HL3_ mutant exhibits higher affinity for Pac1 than wild-type dynein^31^. If this increased affinity is a consequence of Pac1 preferentially binding to an ‘open’, uninhibited dynein conformation, then we reasoned that a disrupted phi particle mutant would also exhibit higher affinity for Pac1 than wild-type dynein. To test this, we assessed the degree of dynein-Pac1 binding using a co-pelleting assay. Either wild-type or dynein^D2868K^ was incubated with Pac1-FLAG-SNAP-decorated beads, and the bound and unbound fractions were quantitatively compared. As a control, we also included GST-dynein_MOTOR_, which we expected to exhibit high Pac1 affinity^3, 48^. Consistent with the notion that Pac1 preferentially binds to dynein in its ‘open’, uninhibited conformation, we found that dynein^D2868K^ and GST-dynein_MOTOR_ both exhibited higher affinity for Pac1 than wild-type dynein (Fig. 4C). This also indicates that the likely reason for the altered localization^31^ and single molecule behavior of dynein^HL3^ is that it is in a constitutively uninhibited state.

### Disruption of the phi particle leads to Pac1-independent cortical dynein activity

Given the phenotypic similarities between the phi particle disrupting mutants and Dyn1_HL3_, we wondered if the Pac1-independent cortical localization of Dyn1_HL3_ is also a property of the phi-disrupting mutants. Specifically, we previously found that the HL3 peptide insertion permits the dynein complex to bypass the need for Pac1 for delivery to cortical Num1 receptor sites^31^. To determine the role of Pac1 in localizing phi particle disrupting mutants to various sites in cells, we imaged Dyn1-3GFP (wild-type or mutant) in cells lacking Pac1 (*pac1*Δ). Consistent with prior observations^44, 48^, Pac1 was required for normal plus end and cortical localization of wild-type dynein (Fig. 4D). This is consistent with the offloading model for dynein function, in which dynein first associates with microtubule plus ends, from where it is delivered – or offloaded – to cortical Num1 sites^31, 49^. Surprisingly, both Dyn1^K1475E^ and Dyn1^D2868K^ were capable of localizing to the cell cortex in the absence of Pac1 (Fig. 4D). The frequency of Dyn1^D2868K^ cortical localization in *pac1*Δ (50%) cells was similar to that of Dyn1_HL3_ (as noted previously^31^, ∼45% of cells exhibit cortical Dyn1_HL3_ foci in *pac1*Δ cells).

We wondered if these Pac1-independent cortical pools of dynein were functional. To assess this, we performed a highly sensitive *in vivo* activity assessment, in which dynein-mediated spindle movements are visualized and quantitated^5^. To eliminate any dynein-independent contributions to spindle movements, these assays were performed in cells lacking Kar9, a protein that is required for an actin/myosin-mediated spindle orientation pathway^50–52^. Generation of yeast strains lacking Kar9 and dynein pathway components also permits a genetic assessment of dynein functionality. In particular, whereas cells deleted for either dynein or Kar9 pathway components exhibit no apparent growth phenotypes, combined deletion of any of the genes involved in these two pathways results in significant synthetic growth defects^53^. As shown in Figure 4E, combined deletion of Kar9 and Pac1 (*kar9*Δ *pac1*Δ) in cells expressing wild-type dynein leads to severe growth defects. Interestingly, *pac1*Δ *kar9*Δ cells expressing Dyn1^D2868K^ exhibited growth defects less severe than those expressing wild-type dynein, suggesting that the D2868K mutation partly rescues the loss of Pac1 (Fig. 4E and Fig. S3A). Interestingly, we did not note a similar rescue for cells expressing Dyn1^HL3^ (Fig. S3B), suggesting that, although this mutant can bypass Pac1 for cortical localization^31^, and it is a highly processive motor *in vitro*, it is unable to move the mitotic spindle in cells.

Consistent with the need for Pac1 for cortical localization of wild-type dynein, we observed no dynein-mediated spindle movements in *pac1*Δ *kar9*Δ cells (Fig. 4F). As expected from the synthetic growth defects in *dyn1^HL3^ kar9*Δ cells, we also observed no dynein-mediated spindle movements in these cells. Surprisingly, we did in fact observe dynein-mediated spindle movements in *dyn1^D2868K^ kar9*Δ *pac1*Δ cells, indicating that the Pac1-independent cortical populations of the uninhibited dynein mutant are indeed active (Fig. 4F and G). Given the ability of the uninhibited dynein mutant to rescue the loss of Pac1, this suggests that at least one function of Pac1 is to release dynein from its autoinhibited phi particle conformation.

### Pac1 stabilizes the uninhibited conformation of motile dynein complexes

To gain additional insight into the potential mechanism by which Pac1 may be affecting dynein autoinhibition we studied available structural information. Specifically, we focused on two structures: a monomeric yeast dynein motor domain bound to Pac1^30^ (with Pac1 bound to a conserved site on the dynein motor domain^26^), and the human dynein complex in the autoinhibited state^7^. Docking of the Pac1-bound motor domain into one of the two motors in the autoinhibited state revealed an apparent steric clash between Pac1 and the motor domain to which Pac1 is not bound (Fig. 5A). This strongly suggests that a Pac1-bound dynein would be precluded from adopting the autoinhibited conformation. This also explains the enhanced affinity of Pac1 for the uninhibited dynein conformation, and also for a truncated dynein motor domain fragment (see Fig. 4C).

**Figure 5.**
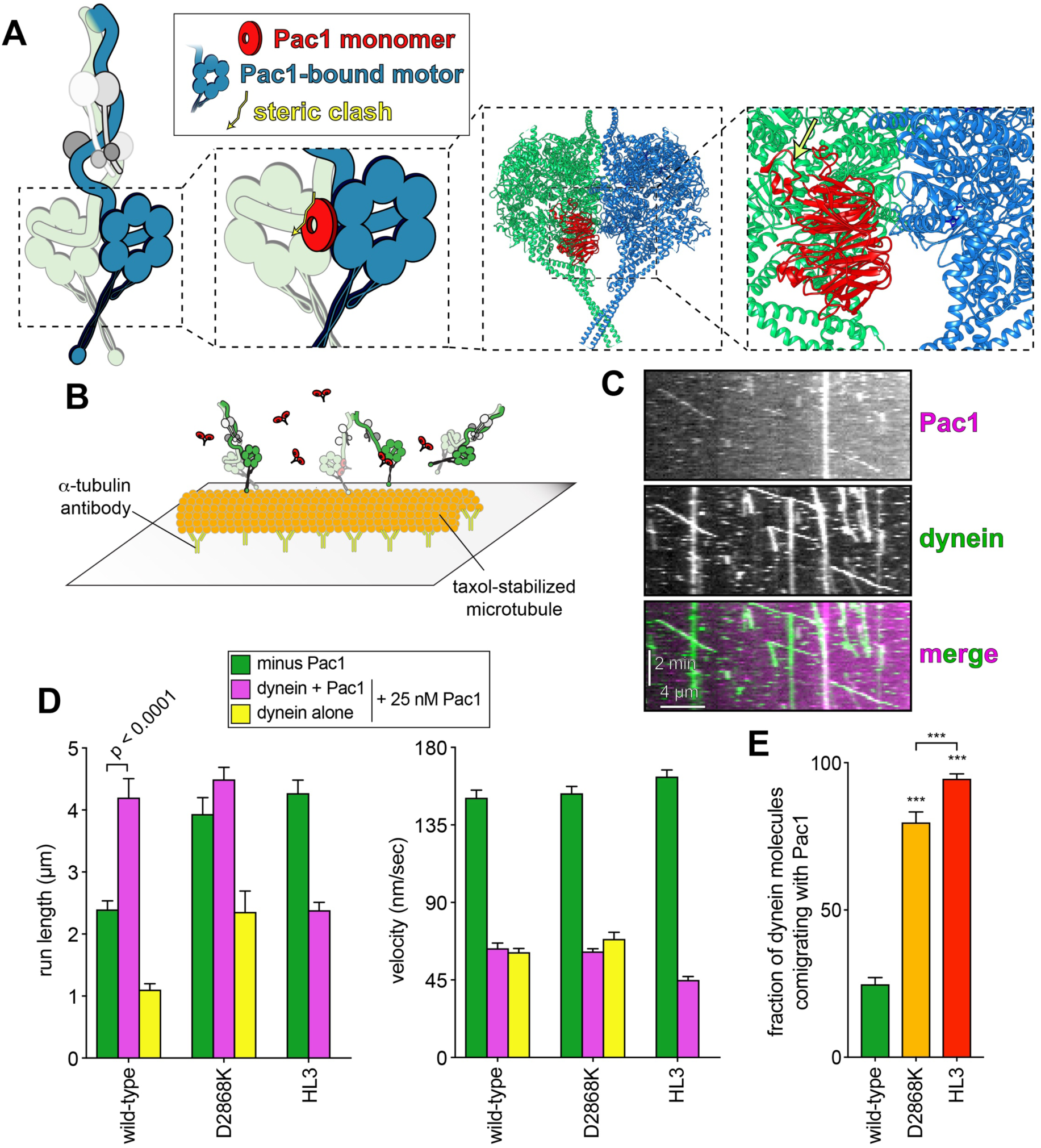
Pac1 promotes release of the autoinhibited conformation of dynein. (A) Cartoon and structural model depicting steric clash between phi particle dynein and Pac1. Structural model was generated by aligning the Pac1-bound dynein monomer structure (pdb 5VH9^30^) into one of the heavy chains in the phi particle structure (pdb 5NVU^7^). Note the steric clash (depicted with jagged yellow arrow) between the Pac1-bound dynein heavy chain (in blue) with the second heavy chain (in green). (B) Cartoon depicting experimental setup for dynein-Pac1 single molecule assay. (C) Representative kymograph illustrating comigrating dynein-Pac1 complexes in motility buffer supplemented with 150 mM potassium acetate (final concentration). (D) Plots depicting motility parameters (left, run length, from fitting of raw data to one-phase decay; right, velocity) of indicated dyneins moving in the absence (*i.e.*, those not pre-incubated with Pac1, green) or presence of 25 nM Pac1 (dimer concentration). For those experiments in which Pac1 and dynein were pre-incubated, we separately scored those dyneins comigrating with Pac1 (magenta), or migrating without Pac1 (yellow; between 134 - 664 dynein ± Pac1 from at least two independent experiments were analyzed for each). To acquire movies of dynein alone, 1-second durations were used; however, for two-color dynein + Pac1 movies, we used 3 second durations due to the speed limitations of our microscope hardware. Statistical significance was determined using a Mann-Whitney test. (E) The fraction of dynein molecules migrating with Pac1 is plotted for the indicated dynein. Error bars depict standard error of proportion. Statistical significance was determined by calculating Z scores (unless indicated by brackets, asterisks indicate statistical difference from wild-type; ***, p < 0.0001; **, p = 0.0011).

To directly test whether Pac1 could affect the conformational state of dynein, we sought to reassess the effect of Pac1 on dynein motility. In light of our single molecule motility data, we predicted that if Pac1 could promote release of dynein autoinhibition, then it would increase dynein run length. Previous studies describing the effect of Pac1 on dynein motility^28, 29, 31^ directly contrast with recent studies of human LIS1^25–27^ (the human homolog of Pac1). Specifically, whereas Pac1 was shown to reduce dynein velocity and promote a microtubule-bound state^28–30^, LIS1 was shown to either increase the velocity of human dynein-dynactin complexes^25, 26^, or have no effect on velocity^27^. In all these studies, LIS1 was observed comigrating with dynein-dynactin complexes at varying degrees. Thus, to clearly define how Pac1 affects dynein motility, we sought to specifically assess comigrating dynein-Pac1 complexes. However, unlike human LIS1^25–27^, we noted that even at nanomolar concentrations, Pac1 bound extensively along microtubules in our motility buffer (Fig. S4A; with 50 mM potassium acetate). We found that the Pac1-microtubule interaction was sensitive to ionic strength: whereas Pac1 strongly bound microtubules in motility buffer supplemented 50 mM potassium acetate, it bound to a much lesser extent in 150 mM potassium acetate (Fig. S4A and B). Thus, we used these latter conditions to assess what effect Pac1 has on dynein motility (Fig. 5B).

Two-color imaging of full-length, wild-type dynein preincubated with Pac1 in motility buffer supplemented with 150 mM potassium acetate revealed many instances of their comigration (Fig. 5C and E). From these movies, we separately scored the run length and velocity values of those dynein molecules that comigrated with Pac1 (Fig. 5D, magenta bars, “dynein + Pac1”), and those that moved without any apparent Pac1 molecules (yellow bars, “dynein alone”). We noted that comigrating dynein-Pac1 complexes moved with significantly longer run lengths than those dynein molecules that were not preincubated with Pac1 (4.2 µm versus 2.4 µm; p < 0.0001). The dynein-Pac1 complexes also moved further than those dynein molecules that were not observed comigrating with Pac1 in the same imaging chamber (4.2 µm versus 1.1 µm). Thus, Pac1 indeed promotes an uninhibited conformational state of motile dynein complexes.

If Pac1 increases run lengths of dynein as a consequence of it promoting an uninhibited state, then we reasoned that Pac1 would not do the same to uninhibited dynein mutants. Consistent with the enhanced affinity of Pac1 for these mutants (Fig. 4C), we observed a much greater frequency of Pac1 molecules comigrating with dynein^D2868K^ and dynein^HL3^ (Fig. 5E). However, we noted no Pac1-dependent increase in run length for either of these mutants, further supporting the notion that are already uninhibited. Note that the mean run length value for wild-type dynein with Pac1 (4.2 µm) is almost identical to that for dynein^D2868K^ with Pac1 (4.5 µm; p = 0.6187), and dynein^HL3^ alone (4.3 µm; p = 0.0620), indicating these all represent similar degrees of uninhibited conformational states. As a side note, consistent with the labile nature of the phi particle conformation of the overexpressed wild-type dynein complex, we noted that the extent of its colocalization with Pac1 increased substantially over time (Fig. S2C and D). Taken together, these data indicate that Pac1 stabilizes the uninhibited conformational state of motile dynein complexes.

### Microtubule-bound Pac1 but not dynein-bound Pac1 reduces dynein velocity

Given the wealth of information pertaining to the Pac1-dependent velocity reduction of dynein^28, 29, 31^, we sought to further address the underlying basis for this phenomenon. Given its propensity to bind microtubules in our motility buffer, we hypothesized that the Pac1-dependent reduction in dynein velocity is a direct consequence of its ability to bind microtubules (in a manner analogous to dynein velocity reduction by the microtubule-associated protein, She1^54, 55^). Consistent with this possibility, structural analysis revealed that Pac1 contacts dynein at a region that is proximal to the microtubule surface (≤ 6.8 nm; Fig. S5A and B; this does not take into account the unstructured E-hooks, which extend away from the surface). Our first clue that this may be the case came from separately analyzing dynein complexes that comigrated with Pac1 versus those that did not (Fig. 5D, yellow versus magenta bars). Given the low, but detectable prevalence of Pac1 along microtubules in these conditions (see Fig. S4A and B), motile dynein complexes still encounter microtubule-bound Pac1 during a processive run. We found that those dynein complexes that comigrated with Pac1 moved with a very similar reduction in velocity as those that did not (Fig. 5D; p = 0.5093). Notably, those dynein complexes that did not comigrate with Pac1 exhibited run lengths somewhat lower that dynein molecules that moved in the absence of Pac1 (compare green and yellow bars). Thus, dynein velocity reduction occurs in a manner that is independent of being bound to Pac1 during a processive run, while processivity enhancement occurs in a manner that requires a stable interaction with Pac1.

To further investigate the effect of Pac1 on dynein velocity, we sought to establish conditions in which Pac1-microtubule binding was further minimized. We found that motility buffer supplemented with 150 mM potassium chloride resulted in a much lower, although still somewhat detectable degree of Pac1-microtubule binding (Fig. S4A and B). Given the sensitivity of dynein microtubule binding and motility to high salt buffers^56^, we chose these conditions as an upper limit for ionic strength for our motility buffer. Importantly, when compared to buffer supplemented with 150 mM potassium acetate, buffer with 150 mM potassium chloride did not appear to negatively impact the Pac1-dynein interaction, as assessed from two-color imaging of dynein^D2868K^ and Pac1 (Fig. 6A).

**Figure 6.**
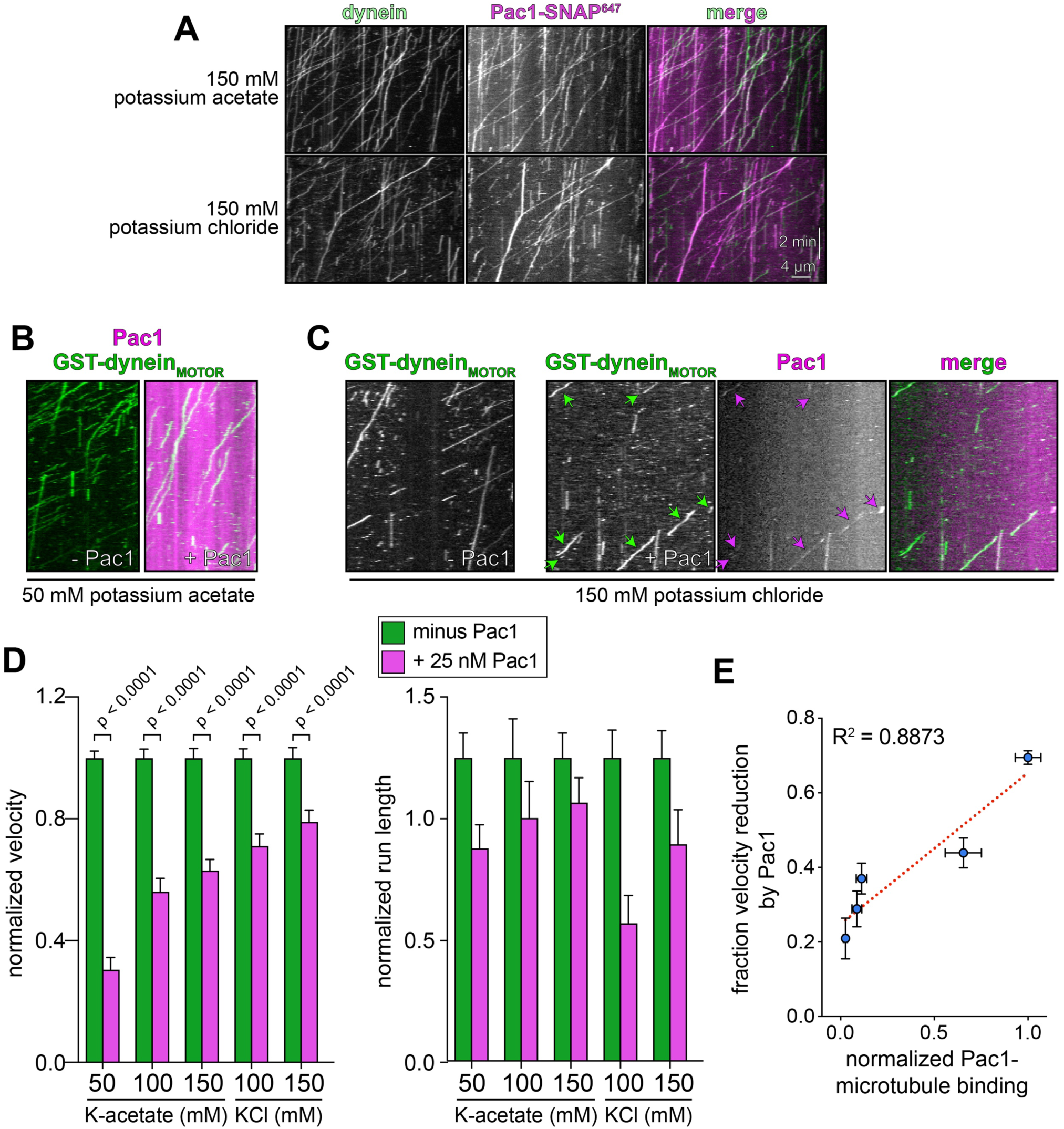
Reducing Pac1-microtubule binding minimizes Pac1-mediated dynein velocity reduction. (A) Representative kymographs depicting dynein^D2868K^ and Pac1 comigrating in single molecule assay in buffers with increasing ionic strength, as indicated. Note that Pac1 and dynein still interact robustly in this assay in both buffer conditions (as apparent by a high degree of colocalization). (B and C) Representative kymographs depicting different motility characteristics of GST-dynein_MOTOR_ in the presence of Pac1 when the latter is either extensively bound to the microtubule (B, in buffer supplemented with 50 mM potassium acetate), or to a much less extent (C, in buffer supplemented with 150 mM potassium chloride). (D) Plots depicting normalized motility parameters (left, normalized run length, from fitting of raw data to one-phase decay; right, normalized velocity) of GST-dynein_MOTOR_ moving in the absence (green) or presence (magenta) of 25 nM Pac1 (dimer concentration; between 226 - 396 motors from two independent experiments were analyzed for each point). Error bars indicate standard error. (E) Plot depicting the relative degree of microtubule binding (normalized to 1; see Fig. S4B) versus the fraction velocity reduction of GST-dynein_MOTOR_ by Pac1. The points (with error bars representing standard error) were fit to a linear regression that indicates a strong correlation between degree of Pac1-microtubule binding and Pac1-mediated dynein velocity reduction.

We next sought to correlate the degree of Pac1-mediated dynein velocity reduction to the extent of microtubule binding by Pac1. We used the GST-dynein_MOTOR_ fragment which has been used extensively in prior Pac1 studies, and exhibits a strong velocity reduction phenotype in low ionic strength buffers^28–30^. As previously reported, 25 nM Pac1 led to a strong (69.5%) reduction in velocity in the low ionic strength buffer (50 mM potassium acetate; Fig. 6B and D, and Fig. S5C and D), in which Pac1 extensively binds along the microtubule lattice (Fig. S4A and B). However, as ionic strength was increased with either potassium acetate or potassium chloride, we noted the effect of Pac1 on GST-dynein_MOTOR_ velocity was substantially reduced (Fig. 6C and D, and Fig. S4D, S5C and D). We plotted the degree of velocity reduction by Pac1 (as shown in Fig. 6D) against the extent of Pac1-microtubule binding in each condition (as shown in Fig. S4B). Fitting these points to a linear regression revealed a very strong correlation between microtubule binding by Pac1 and its ability to affect dynein velocity (Fig. 6E; R^2^ = 0.8873). Thus, in contrast to processivity enhancement by Pac1 – which occurs in a manner that is independent of Pac1-microtubule binding – velocity reduction of dynein by Pac1 appears to occur only when Pac1 is bound to microtubules. Taken together, these findings support a model in which Pac1 is not in fact an inhibitor, but rather an activator of dynein (see Discussion).

## DISCUSSION

In summary, we have shown that like its human orthologue, yeast cytoplasmic dynein adopts an autoinhibited conformation. Furthermore, we have identified a clear biological relevance for this autoinhibited state. Specifically, the phi particle conformation plays a role in coordinating dynein localization and activity within the cell. This becomes abundantly clear in cells expressing uninhibited dynein mutants, which localize to microtubule plus ends and the cell cortex to a greater extent (the former due to an enhanced affinity for Pac1, and the latter due to increased dynactin binding). Moreover, our work identifies a novel role for Pac1 in regulating dynein autoinhibition. Our biochemical, single molecule, genetic, and cell biological data supported by structural analysis reveals the mechanism by which Pac1 modulates the equilibrium between the inhibited and uninhibited states of dynein (see below). Finally, our findings reveal that prior studies describing the role of Pac1 in effecting dynein velocity reduction are for the most part, if not entirely, a consequence of Pac1’s ability to bind microtubules in low ionic strength buffers.

Based on our findings, we propose the following model for dynein function (see Figure 7): (1) Dynein stochastically switches between the inhibited and uninhibited conformational states. (2) Pac1 binds to dynein when it is in the uninhibited state, which consequently prevents dynein from switching to the autoinhibited conformation. (3) The Pac1-dynein complex associates with microtubule plus ends^46^ due to their affinity for Bik1^57^ (homolog of human CLIP-170), which associates with plus ends due in part to its interaction with Bim1^58^ (homolog of human EB1). (4) As a consequence of it being in an uninhibited conformational state^7^, plus end-bound dynein interacts with dynactin. This interaction takes place in the absence of the presumed adaptor molecule, Num1^47^. Although dynein is likely in an open, uninhibited state at microtubule plus ends (due to the requisite presence of Pac1^59^), the precise configuration of the motor domains of this adaptor-free dynein-dynactin complex (which also occurs with human proteins^25, 27, 60^) is unclear. However, the fact that these complexes do not engage in minus end-directed motility – in either budding yeast^47^, or with reconstituted human proteins^25, 27, 60^ – suggests that dynein is inactive at this site, despite being uninhibited. This lack of motility could be due in part to its strong affinity for proteins directly bound to the plus end, and/or due to the dynein heads not being appropriately arranged for proper motility (*i.e.*, in a parallel configuration), which has been observed for human dynein in complex with dynactin and the adaptor BicD2^7^ (see below). (5) Upon encountering Num1 at the cell cortex, the dynein-dynactin complex is offloaded^31^ and activated for motility^47^, possibly due to the arrangement of the motor heads in a parallel manner that is conducive for motility^7^. It is interesting to note that the HL3 mutant, which is our most processive motor in single molecule experiments (see Fig. 4B), is completely inactive in cells, as indicated by our cell biological data (see Fig. 4F). This could be a consequence of the helical linker disrupting the adoption of a parallel head configuration that is potentially needed for cellular dynein-dynactin activity^7^. It also indicates that *in vitro* single molecule motility is not necessarily a good predictor of cellular activity.

**Figure 7.**
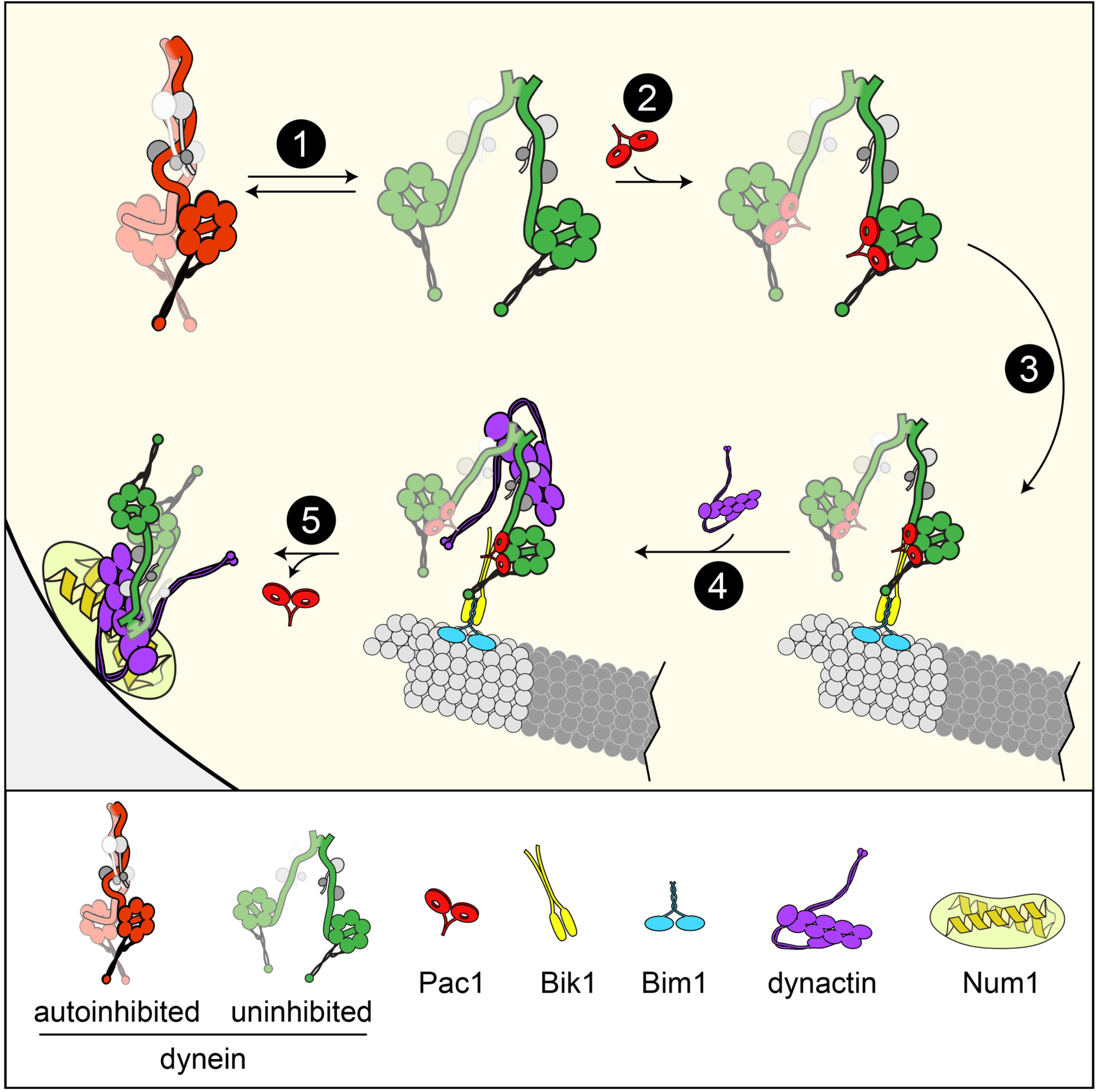
Model for dynein and Pac1 activity in cells. Our data support a model whereby dynein stochastically switches between open and closed states (step 1), the former of which is stabilized by Pac1 binding (step 2). The dynein-Pac1 complex associates with microtubule plus ends via direct interactions with Bik1 (step 3), which associates with plus ends by an unknown mechanism that may rely partly on Bim1. The plus end dynein-Pac1 complex associates with dynactin (step 4), which is then offloaded to cortical Num1 receptor sites (step 5). Given the lack of apparent Pac1 cortical foci, Pac1 likely dissociates either concomitant with, or subsequent to dynein-dynactin offloading.

Rather than acting as an inhibitor, our studies indicate that Pac1 in fact promotes cellular dynein activity by stabilizing the uninhibited conformational state (Fig. 7, step 2). Our data show that prior observations of a Pac1-mediated velocity reduction phenotype^28–31^ are a consequence of the low ionic strength buffers used for these assays. Although we still observe a small effect of Pac1 on dynein velocity even at higher ionic strengths (21.0% reduction), this is likely due to residual microtubule-binding by Pac1 in the highest ionic strength buffer (see Fig. S4A, B and D). These data challenge the current model for Pac1 activity, whereby its binding to the motor domain sterically blocks dynein’s mechanochemical cycle^29^. Our data indicate that Pac1 likely reduces dynein velocity in low ionic strength buffers *in vitro* by simply exerting drag on dynein via simultaneous contacts with dynein and the E-hooks of microtubules (see Fig. S4C and Fig. S5A and B), similar to prior observations with She1 on dynein motility^54^. Given the microtubule-binding-dependent effect of Pac1 on dynein velocity reduction, this raises the question of whether microtubule-binding by Pac1 is a relevant activity in live cells. Several lines of evidence indicate that this is likely not the case. Imaging of Pac1 in live cells reveals it only localizes to microtubule plus ends, and not along the microtubule lattice^44, 46, 61^. The interaction of Pac1 with microtubule plus ends is indirect (as depicted in Figure 7), as it relies on the presence of dynein^46^ and the CLIP-170 homolog, Bik1^46, 61^. Finally, previous studies assessing the role of LIS1 in human dynein function have observed no microtubule-binding activity of LIS1^25–27^. In fact, in contrast to an inhibitory function, two of these studies observed a velocity increase in dynein-dynactin-BicD2 motility due to LIS1^25, 26^. Finally, a previous model describes Pac1 inhibiting dynein release from microtubules as the mechanism by which it prevents dynein from walking away from the plus end^28–30^; however, in direct contradiction to this mode of action, the microtubule-binding domain of dynein is dispensable for its accumulation at microtubule plus ends^47^. In summary, we favor the hypothesis that Pac1 is an activator, not an inhibitor of dynein motility. This role is also supported by evidence in which overexpression of Pac1 leads to increased plus end and cortical localization of dynein, and increased cellular dynein activity^46^.

Although Pac1 is required for dynein localization and activity in cells expressing wild-type dynein, we show that this need for Pac1 can be bypassed when dynein autoinhibition is prevented by mutagenesis (*e.g.*, K1475E and D2868K). This is also apparent by a partial rescue of synthetic growth defects in *kar9*Δ *pac1*Δ cells (see Fig. 4E and Fig. S3A). This is additional support for a role for Pac1 in promoting an uninhibited dynein conformation. It is interesting to note that in spite of the high degree of cortical localization of the uninhibited dynein mutants in *pac1*Δ cells, these cells exhibit much fewer dynein-mediated spindle movements than wild-type cells (Fig. 4F). Thus, offloading of dynein to the cell cortex is more conducive to dynein activity than simple recruitment from the cytoplasm. Although the reasons for this are unclear, we propose that the offloading mechanism is optimally suited to maximize cortical dynein activity. Given the large surface area of the cell cortex (28 µm^2^; assuming a sphere with a 3 µm diameter) with respect to the small number of diffraction limited cortical dynein foci (∼0-2 foci in the daughter cell; *e.g.*, Fig. 3C), the probability of one of the small number of astral microtubules (∼1-2 in the daughter cell) contacting a cortical dynein-dynactin complex to initiate a spindle movement into the daughter cell is very low. However, if the microtubule delivers the motor that will subsequently transport it, the reliance on a stochastic search-based process to initiate a spindle movement event is eliminated. Analysis of dynein-mediated spindle movements in budding yeast revealed that approximately 40% of microtubule-cortex interactions lead to dynein-mediated spindle movements^5^, much greater than would be expected if the cell relied on a stochastic search-based process. Thus, by preventing direct recruitment from the cytoplasm, and requiring a plus end-mediated delivery mechanism, the phi particle, with support from Pac1, ensures that dynein-mediated spindle movements occur within a reasonable timeframe.

In addition to the phi particle restricting localization in cells, we also found that it plays a role in reducing processivity in an *in vitro* context. Given the high proportion of phi particles observed in our negative stain EM images (52.9% in phi conformation, versus 20.0% in an open state; Fig. S1), it is surprising that yeast dynein is processive at all. Our data indicate that yeast dynein stochastically switches from open to closed states in the middle of a processive run. Given the increased processivity of the uninhibited mutants, switching to a closed, inhibited state likely leads to termination of a run, which is likely due to a microtubule release event. This is consistent with the phi particle exhibiting lower affinity for microtubules than the open, uninhibited state^7^. Our findings also raise interesting questions regarding the distinct motility capabilities of yeast versus human dynein, the latter of which requires dynactin and an adaptor for processive single molecule motility^1, 2^. In particular, why is yeast dynein processive without such factors, while human dynein is not? Cryo-EM data of human dynein in the absence and presence of dynactin revealed that dynactin binding orients the motor domains in a parallel configuration, suggesting this is the key to dynactin-triggered processive motility^7^. It is possible that the motor domains of yeast dynein have a higher propensity to adopt a parallel configuration in the absence of dynactin binding; however, higher resolution structural data will be required to determine if this is indeed the case.

Although our work demonstrates a role for the phi particle in budding yeast dynein function, several lines of evidence indicate its importance in humans. For instance, a previous study showed that mutations that disrupt the human dynein phi particle lead to defects in dynein localization and function in human cells^7^. Similar to our observation that yeast dynein mutants localize to the SPBs to a greater extent, human dynein mutants localize more extensively to the spindle poles^7^. As further evidence for the importance of the autoinhibited conformation, at least three different mutations that map to the phi particle contact surfaces have been identified in patients suffering from neurological disease (*i.e.*, malformations in cortical development): E1518K, R1567Q, and R1603T^62–64^ (equivalent to E1425, K1475, and K1510; see residues with “*” in Fig. 2A). We previously showed that a K1475Q dynein mutant exhibits phenotypes much like those described for K1475E (*i.e.*, increased single molecule run lengths and cortical localization), and leads to compromised dynein function in cells^34^. Thus, the phi particle is an important, highly conserved conformational state that is used by organisms throughout evolution to ensure appropriate dynein activity.

## AUTHOR CONTRIBUTIONS

S.M.M. and M.G.M. designed the study. M.G.M. performed the bulk of the *in vitro* and cellular assays, with some support from S.M.M. and J.M.G. The *in vitro* and cellular data were analyzed by M.G.M. Negative staining, grid preparation and electron microscopy was performed by Garry P. Morgan at the University of Colorado Boulder Electron Microscopy facility. Single particle analysis was performed by S.M.M. with assistance from the EM facility support staff. Plasmids were generated by S.M.M. while yeast strains were generated by M.G.M. and J.M.G. The manuscript was written by S.M.M. with assistance from M.G.M.

## ACKNOWLEDGEMENTS

We are grateful to Samara Reck-Peterson for the 8His-ZZ-Pac1-SNAP expressing yeast strain, and members of the Markus and DeLuca laboratories for valuable discussions. Electron microscopy was done at the University of Colorado, Boulder EM Services Core Facility in the MCDB Department, with the technical assistance of facility staff. This work utilized the RMACC Summit supercomputer, which is supported by the National Science Foundation (awards ACI-1532235 and ACI-1532236), the University of Colorado Boulder and Colorado State University. The RMACC Summit supercomputer is a joint effort of the University of Colorado Boulder and Colorado State University. We are also extremely grateful to Erin Osborne-Nishimura, David King, and Samuel Bowerman for their invaluable assistance with using software on Summit. This work was funded by the NIH/NIGMS (GM118492 to S.M.M.).

## METHODS

### Media and strain construction

Strains are derived from either W303 or YEF473A^65^ and are available upon request. We transformed yeast strains using the lithium acetate method^66^. Strains carrying mutations were constructed by PCR product-mediated transformation^67^ or by mating followed by tetrad dissection. Proper tagging and mutagenesis was confirmed by PCR, and in most cases sequencing (all point mutations were confirmed via sequencing). Fluorescent tubulin-expressing yeast strains were generated using plasmids and strategies described previously^68, 69^. Strains overexpressing the yeast dynein complex were generated by transforming p8His-ZZ-SNAPf-Dynein or p8His-ZZ-HALO-Dynein (wild-type or mutants) linearized by digestion with ApaI (cuts within the URA3 gene; see Fig. 1A). Integration was confirmed by PCR. Yeast synthetic defined (SD) media was obtained from Sunrise Science Products (San Diego, CA).

### Plasmid generation

For overexpression and purification of the yeast dynein complex (wild-type or mutants), we generated a polycistronic plasmid expressing all four dynein complex subunits using strategies analogous to the biGBAC assembly^70^. We first made a yeast expression “library” vector – pLIBy – which enables generation of a gene expression cassettes (GEC) with a strong, inducible *GAL1* promoter (*GAL1p*) on the 5’ end, and a synthetic terminator sequence (T_synth3_^37^) on the 3’ end. A PCR product encompassing *GAL1p*, and an oligonucleotide containing T_synth3_^37^ and a multicloning site (XbaI-NotI-SpeI-BamHI) were assembled into pRS305 digested with BamHI and NotI using Gibson assembly^71^, yielding pLIBy. We also generated a yeast genomic-integration vector with optimized linker sequences for Gibson assembly^70^ flanked by PmeI restriction sites (equivalent to pbiG1a and pbiG1b). These plasmids – pbiG1ay and pbiG1by– were generated by using Gibson assembly to insert a PCR product encompassing these elements from pbiG1a and pbiG1b^70^ into pRS306. PCR products encompassing the *DYN2* (without the native intron), *DYN3* or *PAC11* open reading frames were assembled into pLIBy digested with BamHI and NotI. Subsequently, these GECs were amplified from each respective pLIBy vector using oligonucleotides that include regions for priming preceded on the 5’ end by predefined “Cas” sequences^70^: the *DYN2* GEC was amplified with Cas*α*-forward and Cas*β*-reverse; the *DYN3* GEC was amplified with Cas*β*-forward and Cas*γ*-reverse; and, the *PAC11* GEC was amplified with Cas*γ*-forward and Cas*ω*-reverse. These three PCR products were assembled into pbiG1by digested with SwaI to generate pbiG1by:*GAL1p:Dyn2::GAL1p:DYN3::GAL1p:PAC11*.

We generated pLIBy:*6His-StrepII-SNAPf-DYN1* using Gibson assembly. However, due to complications generating a PCR product from this vector, we chose to clone everything into this vector. We first substituted the LEU2 expression cassette in the pLIBy backbone with a URA3 marker by assembling a PCR product encompassing the URA3 cassette from pRS306 into pLIBy:*6His-StrepII-SNAPf-DYN1* digested with KasI and AatII, yielding pLIBy:*6His-StrepII-SNAPf-DYN1::URA3*. To enable assembly of the DYN2/DYN3/PAC11 polygene cassette into pLIBy:*6His-StrepII-SNAPf-DYN1::URA3*, we inserted the optimized “B” and “C” linker sequences for Gibson assembly^70^ into this plasmid by assembling a PCR product encompassing “B”-PmeI site-“C” into pLIBy:*6His-StrepII-SNAPf-DYN1::URA3* digested with KpnI and SalI. Subsequent to digestion with PmeI, this plasmid was assembled with the PmeI restriction digest product from pbiG1by:*GAL1p:Dyn2::GAL1p:DYN3::GAL1p:PAC11* (encompassing *GAL1p:Dyn2::GAL1p:DYN3::GAL1p:PAC11*), yielding pLIBy: *GAL1p:Dyn2::GAL1p:DYN3::GAL1p:PAC11::GAL1p:6His-StrepII-SNAPf-Dyn1::URA3*, hereafter referred to as p6His-StrepII-SNAPf-Dynein. Prior to using this plasmid for pilot tests, we decided to swap the 6His-StrepII affinity tag for an 8His-ZZ tag (followed by a tandem TEV protease recognition site). We did this by assembling a PCR product encompassing 8His-ZZ into p6His-StrepII-SNAPf-Dynein digested with AatII and XhoI, yielding p8His-ZZ-SNAPf-Dynein. We replaced the SNAPf tag with a HALO tag using a similar strategy, yielding p8His-ZZ-HALO-Dynein. All mutations were engineered into these plasmids using common strategies.

### Protein purification

Purification of Pac1-FLAG-SNAP was performed as previously described^28^. Purification of yeast dynein (ZZ-TEV-Dyn1-HALO, under the native *DYN1* promoter; or, ZZ-TEV-HALO-(or SNAPf)-Dynein, with all genes under control of the *GAL1p* promoter; or, ZZ-TEV-6His-GFP-3HA-GST-dynein_MOTOR_-HALO, under the control of the *GAL1p* promoter) was performed as previously described with minor modifications used for the overexpressed complex^28, 54^. Briefly, yeast cultures were grown in YPA supplemented with either 2% glucose (for non-overexpressed full-length dynein) or 2% galactose (for the *GAL1p*-inducible strains), harvested, washed with cold water, and then resuspended in a small volume of water. The resuspended cell pellet was drop frozen into liquid nitrogen and then lysed in a coffee grinder (Hamilton Beach). For most purifications (with exception of those used for negative stain/EM imaging) we used the following procedure: after lysis, 0.25 volume of 4X dynein lysis buffer (1X buffer: 30 mM HEPES, pH 7.2, 50 mM potassium acetate, 2 mM magnesium acetate, 0.2 mM EGTA) supplemented with 1 mM DTT, 0.1 mM Mg-ATP, 0.5 mM Pefabloc SC (concentrations for 1X buffer) was added, and the lysate was clarified at 22,000 × g for 20 min. The supernatant was then bound to IgG sepharose 6 fast flow resin (GE) for 1-1.5 hours at 4°C, which was subsequently washed three times in 5 ml lysis buffer, and twice in TEV buffer (50 mM Tris, pH 8.0, 150 mM potassium acetate, 2 mM magnesium acetate, 1 mM EGTA, 0.005% Triton X-100, 10% glycerol, 1 mM DTT, 0.1 mM Mg-ATP, 0.5 mM Pefabloc SC). To fluorescently label the motors for single molecule analyses, the bead-bound protein was incubated with either 6.7 µM HaloTag-AlexaFluor660 or HaloTag-TMR (Promega), or SNAP-Surface Alex Fluor 647 (NEB), as appropriate, for 10-20 minutes at room temperature. The resin was then washed four more times in TEV digest buffer, then incubated in TEV buffer supplemented with TEV protease for 1-1.5 hours at 16°C. Following TEV digest, the beads were pelleted, and the resulting supernatant was aliquoted, flash frozen in liquid nitrogen, and stored at −80°C. Protein preparations used for negative stain/EM imaging were subject to tandem affinity purification. To do so, subsequent to lysis, 0.25 volume of 4X NiNTA dynein lysis buffer (1X buffer: 30 mM HEPES, pH 7.2, 200 mM potassium acetate, 2 mM magnesium acetate, 10% glycerol) supplemented with 1 mM beta-mercaptoethanol, 0.1 mM Mg-ATP, 0.5 mM Pefabloc SC (concentrations for 1X buffer) was added, and the lysate was clarified as above. The supernatant was then bound to NiNTA agarose for 1 hour at 4°C, which was subsequently washed three times in 5 ml NiNTA lysis buffer. The protein was eluted in NiNTA lysis buffer supplemented with 250 mM imidazole by incubation for 10 minutes on ice. The eluate was then diluted with an equal volume of dynein lysis buffer, which was then incubated with IgG sepharose 6 fast flow resin for 1 hour at 4°C. The beads were washed and the protein was eluted as described above. Eluted protein was either applied to a size exclusion resin (Superose 6; GE), or snap frozen. The gel filtration resin was equilibrated in GF150 buffer (25 mM HEPES pH 7.4, 150 mM KCl, 1 mM MgCl_2_, 5 mM DTT, 0.1 mM Mg-ATP) using an AKTA Pure. Peak fractions (determined by UV 260 nm absorbance and SDS-PAGE) were pooled, concentrated, aliquoted, flash frozen, then stored at −80°C.

For comparison of elution profiles between yeast and human dynein complexes, the human dynein complex was expressed and purified from insect cells (ExpiSf9 cells; Life Technologies) as previously described with minor modifications^1, 7^. Briefly, 4 ml of ExpiSf9 cells at 2.5 × 10^6^ cells/ml, which were maintained in ExpiSf CD Medium (Life Technologies), were transfected with 1 µg of bacmid DNA (see above) using ExpiFectamine (Life Technologies) according to the manufacturer’s instructions. 5 days following transfection, the cells were pelleted, and 1 ml of the resulting supernatant (P1) was used to infect 300 ml of ExpiSf9 cells (5 × 10^6^ cells/ml). 72 hours later, the cells were harvested (2000 × g, 20 min), washed with phosphate buffered saline (pH 7.2), pelleted again (1810 × g, 20 min), and resuspended in an equal volume of human dynein lysis buffer (50 mM HEPES, pH 7.4, 100 mM NaCl, 10% glycerol, 1 mM DTT, 0.1 mM Mg-ATP, 1 mM PMSF). The resulting cell suspension was drop frozen in liquid nitrogen and stored at −80°C. For protein purification, 30 ml of additional human dynein lysis buffer supplemented with cOmplete protease inhibitor cocktail (Roche) was added to the frozen cell pellet, which was then rapidly thawed in a 37°C water bath prior to incubation on ice. Cells were lysed in a dounce-type tissue grinder (Wheaton) using ≥ 150 strokes (lysis was monitored by microscopy). Subsequent to clarification at 22,000 × g, 45 min, the supernatant was applied to 2 ml of IgG sepharose fast flow resin pre-equilibrated in human dynein lysis buffer, and incubated at 4°C for 2-4 hours. Beads were then washed with 50 ml of human dynein lysis buffer, and 50 ml of human dynein TEV buffer (50 mM Tris pH 7.4, 150 mM potassium acetate, 2 mM magnesium acetate, 1 mM EGTA, 10% glycerol, 1 mM DTT, 0.1 mM Mg-ATP). The bead-bound protein was eluted with by incubation with TEV protease overnight at 4°C. The next morning, the recovered supernatant was applied to a Superose 6 gel filtration column as above.

### Single molecule motility assays

The yeast dynein single-molecule motility assay was performed as previously described with minor modifications^54^. Briefly, flow chambers constructed using slides and plasma cleaned and silanized coverslips attached with double-sided adhesive tape were coated with anti-tubulin antibody (8 μg/ml, YL1/2; Accurate Chemical & Scientific Corporation) then blocked with 1% Pluronic F-127 (Fisher Scientific). Taxol-stabilized microtubules assembled from unlabeled and fluorescently-labeled porcine tubulin (10:1 ratio; Cytoskeleton) were introduced into the chamber. Following a 5-10 minute incubation, the chamber was washed with dynein lysis buffer (see above) supplemented with 20 μM taxol. Subsequently, purified dynein motors diluted in motility buffer (30 mM HEPES pH 7.2, 2 mM magnesium acetate, 1 mM EGTA, 1 mM DTT, 1 mM Mg-ATP, 0.05% Pluronic F-127, 20 µM taxol, and an oxygen-scavenging system consisting of 1.5% glucose, 1 U/ml glucose oxidase, 125 U/ml catalase) supplemented with either 50 mM potassium acetate, or as indicated in figure legend, were introduced in the chamber, and imaged.

To image comigrating Pac1-dynein complexes, 500 nM Pac1-SNAP^6^^47^ (dimer concentration) and ∼50 nM HALO^TMR^-Dynein were preincubated on ice for 10-15 minutes prior to a 20-fold dilution into modified motility buffer (30 mM HEPES pH 7.2, 2 mM magnesium acetate, 1 mM EGTA, 1 mM DTT, 1 mM Mg-ATP) supplemented with potassium acetate or potassium chloride as indicated in figure legends, 0.05% Pluronic F-127, 20 µM taxol, and an oxygen-scavenging system (as above). The higher yield overexpressed dynein complex was needed for these assays given the low landing rate of dynein in the higher ionic strength buffers. We ensured that comigrating Pac1-SNAP^647^ spots were not due to bleed-through from the HALO^TMR^-dynein channel by performing two-color imaging with HALO^TMR^-dynein alone (no spots were apparent in the far-red channel in these cases).

TIRFM images were collected using a 1.49 NA 100X TIRF objective on a Nikon Ti-E inverted microscope equipped with a Ti-S-E motorized stage, piezo Z-control (Physik Instrumente), and an iXon X3 DU897 cooled EM-CCD camera (Andor). 488 nm, 561 nm, and 640 nm lasers (Coherent) were used along with a multi-pass quad filter cube set (C-TIRF for 405/488/561/638 nm; Chroma) and emission filters mounted in a filter wheel (525/50 nm, 600/50 nm and 700/75 nm; Chroma). We acquired images at 1, 2, or 3 second intervals for 8-10 min. Velocity and run length values were determined from kymographs generated using the MultipleKymograph plugin for ImageJ (http://www.embl.de/eamnet/html/body_kymograph.html). Those motors that moved for ≥ 3 time points were measured.

### Negative stain electron microscopy and image analysis

EM grids were prepared with a standard negative stain protocol by applying fresh dynein samples to glow discharged carbon coated 200 mesh copper grids. After ∼1 minute incubation, 2% uranyl acetate was added. 1600 micrographs were collected on a FEI Tecnai F20 200kV TEM equipped with a Gatan US4000 CCD (model 984), at a nominal magnification of 90,000X with the digital pixel size 6.19 angstroms. All image analysis was performed in Relion 3.0 on the University of Colorado Boulder High Performance Computer Cluster, Summit. Particles were manually picked from ∼20 micrographs (∼200 particles), which were used to generate a low resolution 2D class average. Using these 2D averages as a starting point, we then used an iterative process to autopick particles that were used to generate our final 2D averages, and for 3D model building (in total, 42,611 particles were used for final averages shown in Figure 1D).

### Live cell imaging experiments

For the spindle dynamics assay, cells were arrested with hydroxyurea (HU) for 2.5 hours, and then mounted on agarose pads containing HU for fluorescence microscopy. Full Z-stacks (23 planes with 0.2 µm spacing) of GFP-labeled microtubules (GFP-Tub1) were acquired every 10 seconds for 10 minutes on a stage prewarmed to 30°C. To image dynein localization in live cells, cells were grown to mid-log phase in SD media supplemented with 2% glucose, and mounted on agarose pads. Images were collected on a Nikon Ti-E microscope equipped with a 1.49 NA 100X TIRF objective, a Ti-S-E motorized stage, piezo Z-control (Physik Instrumente), an iXon DU888 cooled EM-CCD camera (Andor), a stage-top incubation system (Okolab), and a spinning disc confocal scanner unit (CSUX1; Yokogawa) with an emission filter wheel (ET480/40m for mTurquoise2, ET525/50M for GFP, and ET632/60M for mRuby2; Chroma). Lasers (445 nm, 488 nm and 561 nm) housed in a LU-NV laser unit equipped with AOTF control (Nikon) were used to excite mTurquoise2, GFP and mRuby2, respectively. The microscope was controlled with NIS Elements software (Nikon).

### Statistical analyses

Statistical tests were performed as described in the figure legends. T-tests were performed using Graphpad Prism. Z scores were calculated using the following formula:

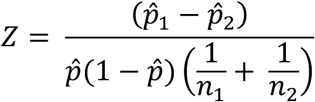

where:

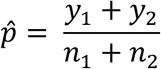

Z scores were converted to two-tailed P values using an online calculator.

### Data availability

All yeast strains, and datasets generated during and/or analysed during the current study are available from the corresponding author upon request.

