## Supplemental Information for "Pac1/LIS1 promotes an uninhibited conformation of dynein that coordinates its localization and activity"

#### Supplementary Figures and Figure Legends

### Figure S1

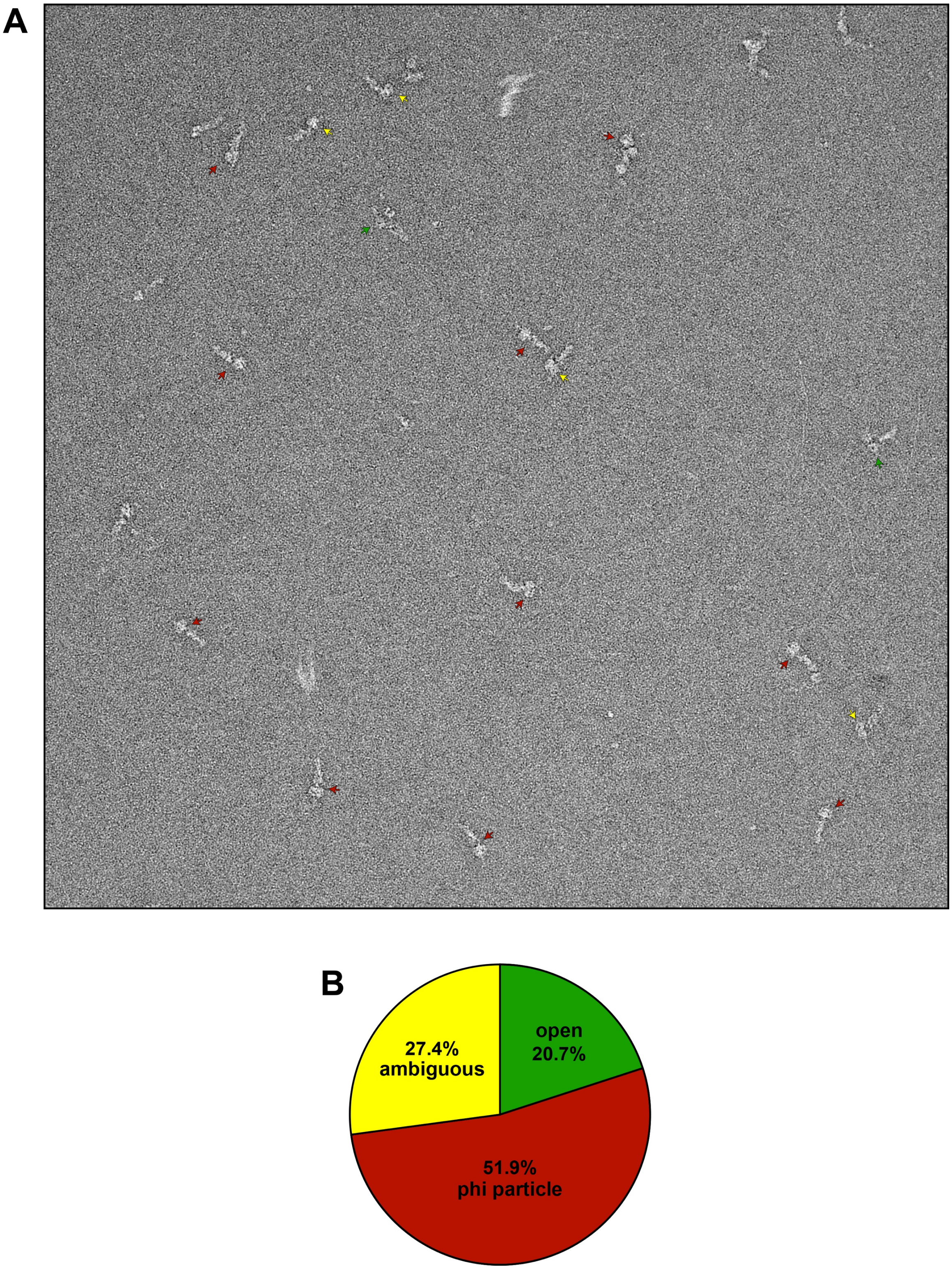

Figure S1. **Representative raw EM image and quantitation of conformational states.** (A) Representative EM image of negative stained yeast dynein complex (red arrow, phi particle conformation; green arrow, open conformation; yellow arrow, ambiguous). (B) Quantitation of various conformational states from raw images (n = 435 particles).

### Figure S2

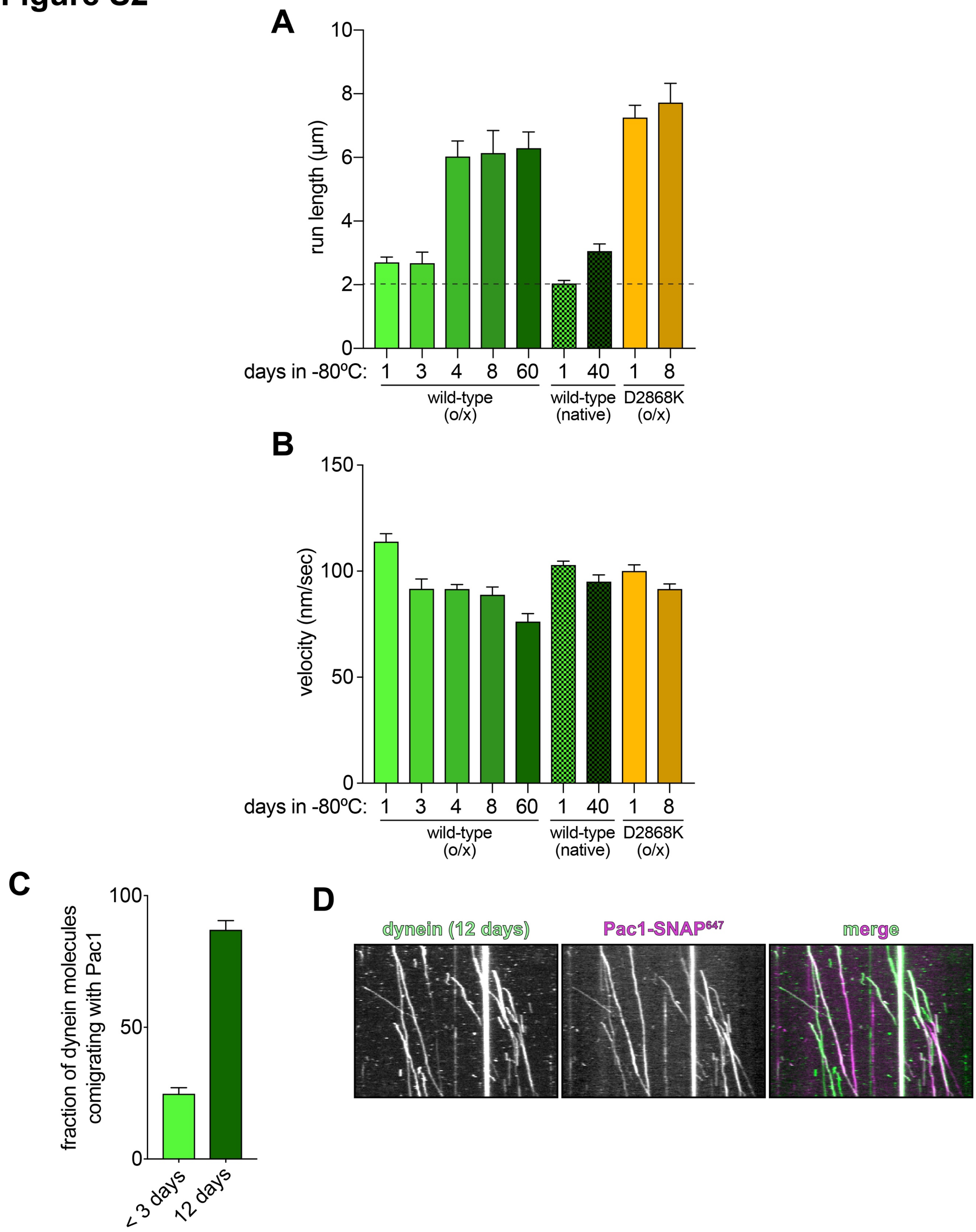

A
run length (µm)
10
8
6
4
2
0
days in -80ºC: 1 3 4 8 60 1 40 1 8
wild-type (o/x)
wild-type (native)
D2868K (o/x)
B
velocity (nm/sec)
150
100
50
0
days in -80ºC: 1 3 4 8 60 1 40 1 8
wild-type (o/x)
wild-type (native)
D2868K (o/x)
C
fraction of dynein molecules comigrating with Pac1
100
50
0
< 3 days
12 days
D
dynein (12 days)
Pac1-SNAP647
merge

Figure S2. **Overexpressed yeast dynein complex exhibits uninhibited properties over time.** (A and B) Plots depicting mean single molecule motility parameters (A, run length from fitting of raw data to one-phase decay; B, velocity) with standard error (between 67 and 527 motors were analyzed for each point; "o/x", over-expressed dynein complex, as described in Figure 1; "native", non-over-expressed dynein complex, as described previously[1]). Proteins were stored at -80°C between time points. Note that run length values for wild-type dynein increase to levels similar to that of dynein$^{D2868K}$, but the latter only increases to a minimal extent over the same time period. Also note that run length values for the non-overexpressed dynein complex (expressed from native promoters, "native") only increased to a minor extent in comparison to the overexpressed complex. (C) Fraction of overexpressed, wild-type dynein complexes comigrating with Pac1 in single molecule assays (in motility buffer supplemented with 150 mM potassium acetate). (D) Representative kymograph of 12 day old, over-expressed dynein comigrating with Pac1. Note the large increase in the frequency of comigrating complexes over time. Taken together, these data indicate that the overexpressed complex transitions to an uninhibited state over time, similar to observations with recombinantly expressed human dynein[2].

### Figure S3

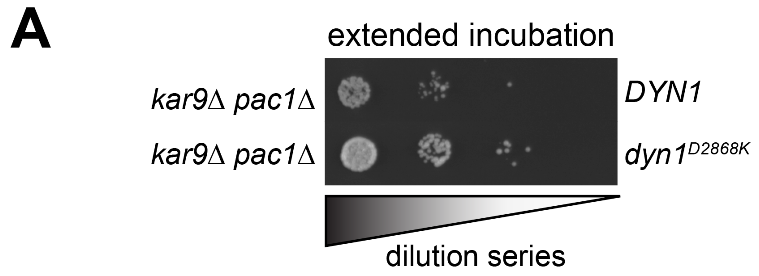
A
extended incubation
kar9Δ pac1Δ
DYN1
kar9Δ pac1Δ
dyn1D2868K
dilution series

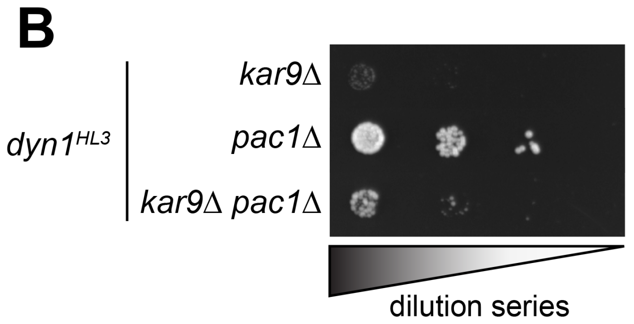
B
dyn1HL3
kar9Δ
pac1Δ
kar9Δ pac1Δ
dilution series

Figure S3. **Synthetic interactions between dynein mutants and Kar9.** (A and B) Serial dilutions of cells with indicated genotype were plated on rich media (YPA supplemented with 2% glucose) and grown at 30°C for 4 days (A; extended incubation of plates from Fig. 4E), or 2 days (B). Note the severe growth defects in $dyn1^{HL3}$ *kar9*Δ cells, suggesting that $Dyn1^{HL3}$ is not active in cells.

### Figure S4

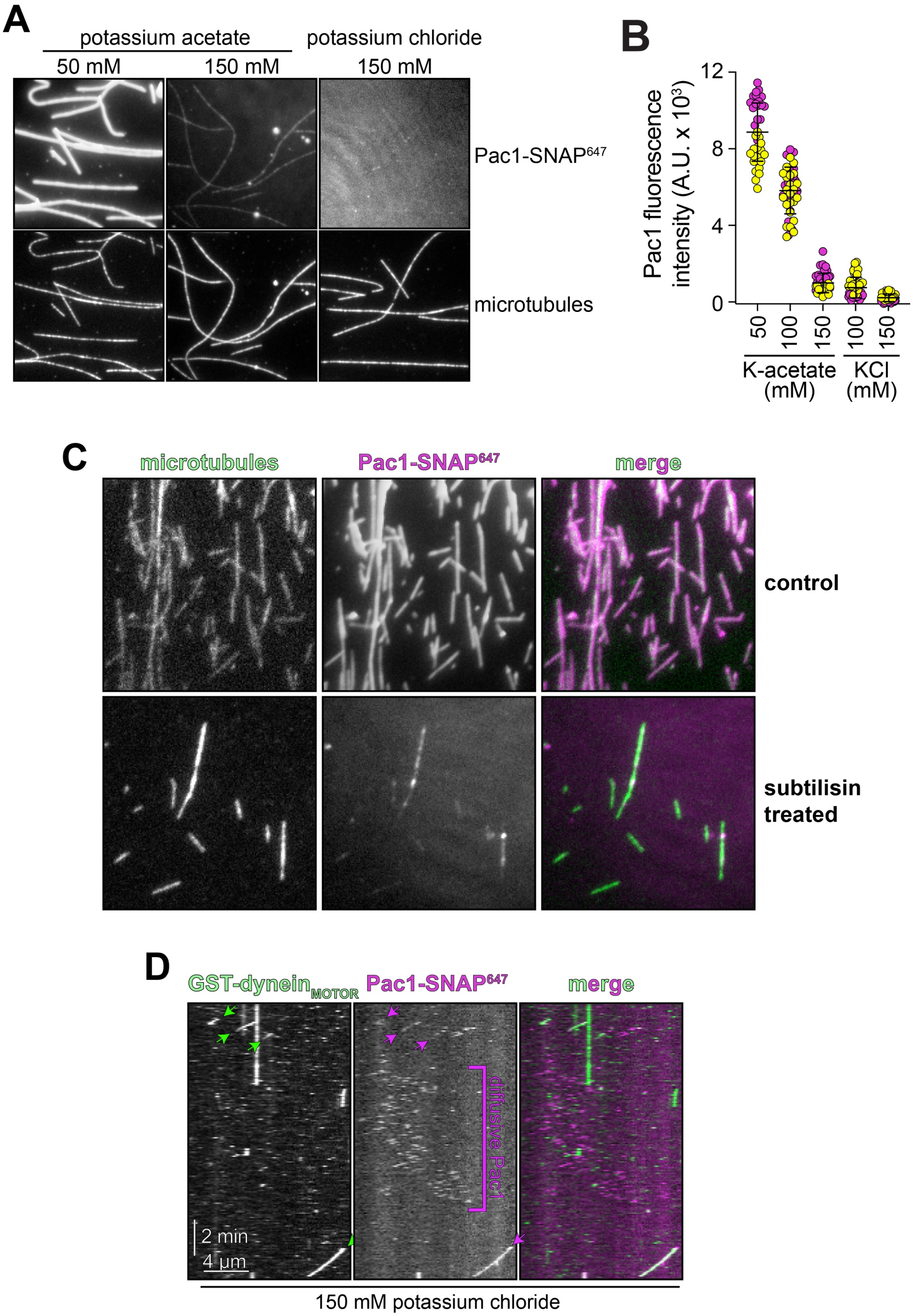
A
potassium acetate
potassium chloride
50 mM
150 mM
150 mM
Pac1-SNAP647
microtubules
B
Pac1 fluorescence intensity (A.U. x 10³)
12
8
4
0
50
100
150
100
150
K-acetate (mM)
KCl (mM)
C
microtubules
Pac1-SNAP647
merge
control
subtilisin treated
D
GST-dyneinMOTOR
Pac1-SNAP647
merge
diffusive Pac1
2 min
4 µm
150 mM potassium chloride

Figure S4. **Characterization of Pac1-microtubule binding behavior.** (A and B) Representative fluorescence images (A) and quantitation (B) of Pac1 binding to microtubules in motility buffer (see methods) supplemented with indicated salts. Pac1-$SNAP^{647}$ was diluted into motility buffer with indicated salt conditions (to 50 nM final, dimer concentration), and images were acquired of microtubules and Pac1 (yellow and magenta circles represent data acquired from two independent experiments; $n \geq 38$ microtubules spanning at least 910 $\mu$m in length for each condition). (C) Pac1-microtubule binding is reduced after enzymatic removal of the unstructured carboxy-terminal tails of $\alpha$-tubulin and $\beta$-tubulin. Briefly, taxol-stabilized microtubules were digested with a freshly dissolved preparation of 1 mg/ml subtilisin (Sigma) for 60 min at 37 °C prior to preparation of flow chambers (see Methods). (D) An additional representative kymograph of GST-$dynein_{MOTOR}$ comigrating with Pac1 in motility buffer supplemented with 150 mM potassium chloride. Note the diffusive behavior of Pac1 on microtubules in this example.

### Figure S5

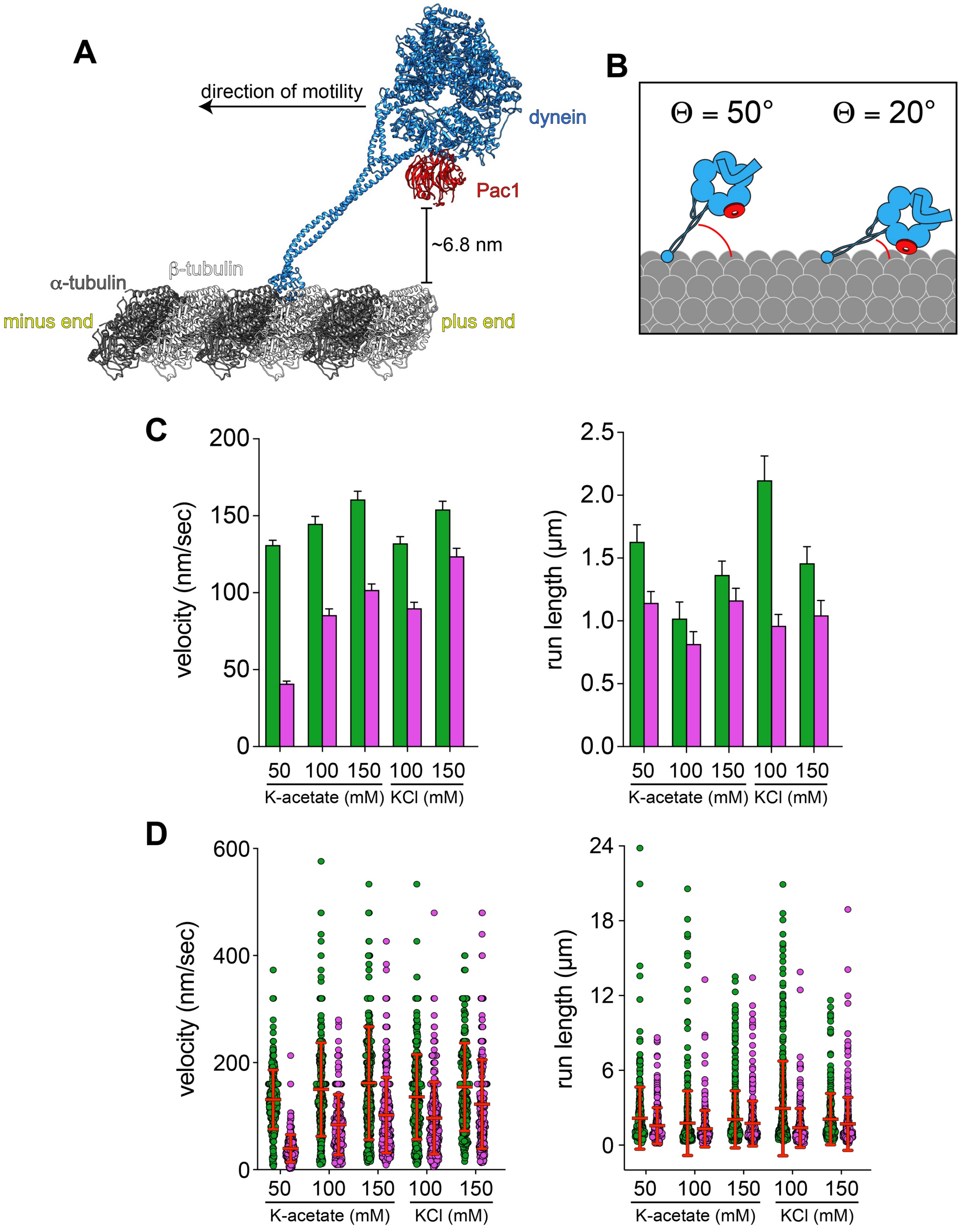
A
direction of motility
dynein
Pac1
~6.8 nm
α-tubulin
β-tubulin
minus end
plus end
B
Θ = 50°
Θ = 20°
C
velocity (nm/sec)
run length (µm)
50 100 150 100 150
K-acetate (mM)
KCl (mM)
D
velocity (nm/sec)
run length (µm)
50 100 150 100 150
K-acetate (mM)
KCl (mM)

Figure S5. **Structural analysis of the Pac1-dynein-microtubule complex, and additional plots of dynein motility.** (A and B) Structural and cartoon model of a Pac1-bound dynein monomer on microtubules. The following structures were used to generate the model shown in panel A: pdb 4RH7[3] (human dynein-2 motor domain), pdb 3J1T[4] (yeast dynein microtubule-binding domain bound to tubulin dimer), pdb 5VH9[5] (yeast dynein monomer bound to Pac1), and pdb 3J6G[6] (taxol-stabilized microtubule). Note the close proximity of Pac1 to the microtubule surface, the latter of which is lacking the unstructured C-terminal tails. (B) Cryo-EM images of microtubule-bound dynein reveals the stalk angle with respect to the microtubule surface varies due to a hinge point within the microtubule-binding domain, and can be much steeper than that shown in panel A[7,8] ($\Theta \geq$ 15-20°, with an average of 55°). Cartoons depicts range of angles sampled by dynein on microtubules, and thus the distances between Pac1 and the microtubule that are sampled. (C and D) Non-normalized plots of mean values (C) and all data points (D) showing the relationship between Pac1-mediated dynein velocity reduction and Pac1-microtubule binding.
